# Common genetic variation indicates separate etiologies for periventricular and deep white matter hyperintensities

**DOI:** 10.1101/683367

**Authors:** Nicola J Armstrong, Karen A Mather, Muralidharan Sargurupremraj, Maria J Knol, Rainer Malik, Claudia L Satizabal, Lisa R Yanek, Wen Wei, Vilmundur Gudnason, Nicole D Deuker, Lloyd T Elliott, Edith Hofer, Neda Jahanshad, Shuo Li, Mark A Logue, Michelle Luciano, Markus Scholz, Albert Smith, Stella S Trompet, Dina Vojinovic, Rui Xia, Fidel Alfaro-Almagro, David Ames, Najaf Amin, Philippe Amouyel, Alexa S Beiser, Henry Brodaty, Ian J Deary, Christine Fennema-Notestine, Piyush G Gampwar, Rebecca Gottesman, Ludovica Griffanti, Clifford R Jack, Mark Jenkinson, Jiyang Jain, Brian G Kral, John W Kwok, Leonie Lampe, David CM Liewald, Pauline Maillard, Jonathan Marchini, Mark E Bastin, Bernard Mazoyer, Lukas Pirpamer, José Rafael Romero, Gennady V Roshchupkin, Peter R Schofield, Matthias L Schroeter, David J Stott, Anbupalam Thalamuth, Julian Trollor, Christophe Tzourio, Jeroen van der Grond, Meike W Vernooij, Veronica A Witte, Maragret J Wright, Qiong Yang, Moris Zoe, Siggi Siggurdsson, Arno Villringer, Helena Schmidt, Asta L Haberg, Cornelia M Van Duijn, J Wouter Jukema, Martin Dichigans, Ralph L Sacco, Clinton B Wright, William S Kremen, Lewis C Becker, Paul M Thompson, Lenore Launer, Thomas H Mosley, Joanna M Wardlaw, M Afran Ikram, Hieab HH Adams, Reinhold Schmidt, Stephen M Smith, Charles Decarli, Perminder S Sachdev, Myriam Fornage, Stephanie Debbette, Sudha Seshadri, Paul A Nyquist

## Abstract

We conducted a genome-wide association meta-analysis of two ischemic white matter disease subtypes in the brain, periventricular and deep white matter hyperintensities (PVWMH and DWMH). In 26,654 participants, we found 10 independent genome-wide significant loci only associated with PVWMH, four of which have not been described previously for total WMH burden (16q24.2, 17q21.31, 10q23.1, 7q36.1). Additionally, in both PVWMH and DWMH we observed the previous association of the 17q25.1 locus with total WMH. We found that both phenotypes have shared but also distinct genetic architectures, consistent with both different underlying and related pathophysiology. PVWMH had more extensive genetic overlap with small vessel ischemic stroke, and unique associations with several loci implicated in ischemic stroke. DWMH were characterized by associations with loci previously implicated in vascular as well as astrocytic and neuronal function. Our study confirms the utility of these phenotypes and identifies new candidate genes associated only with PVWMH.

## Introduction

Radiological white matter hyperintensities of presumed ischemic origin (WMH, white matter lesions, leukoaraiosis) are the most prevalent sign of cerebral small vessel disease (SVD) and represent 40% of all SVD disease burden^1^. WMH begin to be observed in middle age, are present in approximately 50% of adults by their mid to late 40’s and are commonly found in community-dwelling older adults^2^. They are detected as incidental lesions on T2-weighted MRI^1^. They are one of many radiological phenotypes associated with SVD including: microhemorrhages, enlarged perivascular spaces, gliosis, and brain atrophy^1^. WMH are associated with increased risk for ischemic and hemorrhagic stroke, cognitive decline that may result in dementia, and motor gait disorders and reflect ongoing chronic subclinical brain injury^3-7^. They are also commonly related to histopathological changes caused by hypertension and other risk factors such as microvascular occlusion, lipohyalinosis, gliosis, and fibrosis of surrounding adventitia^6,8,9^. These lesion types are important clinical targets for intervention in prevention of cognitive aging, stroke, and dementia^1,10-12^. Total WMH burden has high heritability and several genome-wide association studies (GWAS) have been previously reported^13-19^.

Two important and common regional classifications, based on their anatomical relationship to the lateral ventricles in the brain, are periventricular white matter hyperintensities (PVWMH) and deep white matter hyperintensities (DWMH)^6,8,20-22^. This categorization reflects proposed differences in underlying pathophysiology, histopathology, and etiology for each lesion type^6,8,20,21^. Both lesion types are hypothesized to reflect chronic microvascular injury from hypoperfusion and/or microvascular occlusion^1,6,8^. The microvasculature subserving PVWMH and DWMH differ based on the type of microvessels that supply these brain regions. DWMH lesions occur in the subcortex, areas primarily supplied by long microvessels, with lower estimated blood pressures thought subject to hypoperfusion^1,8,23,24^. In contrast, PVWMH lesions are related to alterations in short penetrating microvessels ending in close approximation to larger arterial blood vessels with different vascular architecture such as two leptomeningeal layers and enlarged perivascular spaces^1,21,25^. These microvessels are more directly perfused by large systemic arterial vessels and have much higher estimated mean arterial pressures than those supplying DWMH. These vessels are hypothesized to be affected more directly by hypertension and other risk factors associated with stroke^1,8,23,24^.

PVWMH and DWMH are associated with different clinical phenotypes. PVWMH are associated with cognitive performance and increased systolic and arterial pressure, while DWMH are linked to BMI, mood disorders, gait impairment and arterial hypertension ^12,26,27^. These sub-classifications may reflect differences in associated underlying genetic factors^13^. Twin and family studies report that both PVWMH and DWMH have high heritability^13,14^. Genetic correlations between the two phenotypes are high^13,28^. Recently, GWAS for total WMH volume identified a major genetic risk locus on chromosome 17q25.1^15,16,18,19^. Several other genome-wide significant loci were identified (e.g., 10q24, 2p21, 2q33, 6q25.1)^16,19,29^. However, the genetic determinants of regional WMH burden, specifically DWMH and PVWMH, remain elusive.

Here we combined all available participants aged 45 and above with both DWMH and PVWMH measurements from the Cohorts for Heart and Aging Research in Genomic Epidemiology (CHARGE) and the Enhancing Neuro-Imaging Genetics through Meta-Analysis (ENIGMA) consortia, and the UK Biobank (UKB). The combined sample size of this study was n=26,654 and is the only GWAS to date of these subclassifications of WMH. We hypothesized that separating the two WMH subclassifications would mitigate phenotype heterogeneity, allowing us to identify additional risk loci and show that DWMH and PVWMH have different genetic underpinnings. We further estimated the genetic correlations of DWMH and PVWMH with stroke, dementia, and vascular disease, hypothesizing that there would be differences between the two subclassifications.

## Methods

### Study Cohorts

Study participants (total N∼26,654) were drawn from studies of the CHARGE and ENIGMA consortia and UK Biobank (UKB). All participants were aged 45 years and over and individuals with stroke, dementia or any neurological abnormality such as tumors were excluded. Descriptions of the individual studies are found in the Supplementary Note. The demographic details of the included participants from each study are presented in Supplementary Table 1. All participants provided written informed consent and each study received ethical approval to undertake this work.

### Phenotype and covariates

The majority of studies (n=14) extracted deep and periventricular WMH volumetric data using automated methods while three studies (HUNT, LBC, AGES, n= 3843) used visual rating scales. Volumetric measures were transformed by adding 1 and taking the natural logarithm. The UKB processed their volumetric WMH data as per Elliott et al^30^. The MRI and WMH extraction methods for each study are detailed in Supplementary Table 2. For all participants from CHARGE and ENIGMA, hypertension was defined as systolic blood pressure ≥140mm Hg and diastolic blood pressure ≥90mm Hg or on current antihypertensive treatment and coded as yes (1) versus no (0). For volumetric measures, intracranial volume (ICV) was included as a covariate unless ICV had already been adjusted for when the WMH measures were calculated.

### Genotyping and Imputation

Genotyping platforms, imputation and quality control procedures used for each participating study are described in Supplementary Table 3. For the majority of studies, genotype data were imputed using the 1000 Genomes Project reference panel (phase 1, version 3) using standard software and methods. One cohort, VETSA, used the phase 3, version 5 panel. The UKB details and procedures are detailed in Elliott et al^30^.

### Statistics

All statistical tests were two-sided and, unless stated otherwise, the conventional *p*-value of 0.05 was used for significance. When necessary, Bonferroni correction for multiple testing was applied.

### Genome-wide association analysis

Each study fitted linear regression models to test the association of DWMH and PVWMH with individual SNPs. Additive genetic effects were assumed and the models were adjusted for age (years), sex and ICV (where applicable). In addition, principal components for population stratification and other covariates, such as familial structure, were included if necessary. Models were also fitted with hypertension as an additional covariate. Details of the covariates included for each study are listed in Supplementary Table 3.

Prior to meta-analysis, quality control of summary statistics for each cohort was performed using EasyQC^31^. Marker names, alleles and chromosome base pair positions were harmonized to the appropriate 1000 Genome reference panel. Meta-analyses were restricted to biallelic SNPs. At the individual study level, SNPs were removed due to duplication, missing information, allelic mismatch to the reference panel or allele frequency outliers (where the study allele frequency differed from the reference frequency by more than 0.2). The effective allele count was defined as twice the product of the MAF, imputation accuracy (r^2^, info score or oevar_imp), and the study sample size. Variants with an effective allele count < 10 were excluded. QQ plots, allele-frequency plots and P-Z plots were constructed for each study and visually checked pre and post QC.

Fixed-effects, inverse-variance-weighted meta-analysis was carried out in METAL^32^, with correction for genomic control. Two meta-analyses were carried out: all cohorts excluding UKB (discovery, phase I) and all cohorts (phase II). Post meta-analysis QC, filtering SNPs by their heterozygosity index (I^2^>0.8) and sample size (phase I: N> 11,000; phase II: N>14,000) was also performed.

SNPs were denoted as genome-wide significant if p<5e-8, and suggestive if p<1e-5. Replication of the phase I genome-wide significant SNPs was performed in the UKB. The *p*-value for replication was set to the nominal significance level (0.05) divided by total number of genome-wide significant SNPs (DWMH *p*< 3.60e-4; PVWMH, *p*<2.75e-4). Manhattan and Q-Q plots for the meta-analysis were created in FUMA^33^. Regional association plots were generated with LocusZoom^34^. Annotation of genome-wide significant SNPs was performed using FUMA^35^. Further *in silico* functional annotation was undertaken for the top SNPs using publicly available databases (SNiPA (http://snipa.helmholtz-muenchen.de/snipa/), SNPnexus (https://www.snp-nexus.org/), and GTEx (http://www.gtexportal.org/home/))^35-37^.

### Gene-based and pathway analysis

Gene-based tests and gene set (pathway) analyses were performed using MAGMA as implemented in FUMA^33,38^. For gene-based tests, significance was set to 2.667e-6 (0.05 divided by the number of protein coding genes (18745)). For biological pathways, 10655 gene sets (4738 curated gene sets and 5917 GO terms from MsigDB v6.1) were tested, resulting in a significance level of 4.69e-6.

### Heritability

In the Rotterdam study, SNP-based heritability was computed with GCTA^39^ adjusting for age and sex. Additionally, univariate genome-wide heritability estimates were calculated using the phase II GWAS summary statistics, for both DWMH and PVWMH, using LD score regression (LDSC)^40^ and the European LD-score files calculated from the 1000G reference panel provided by the developers.

### Genetic correlations with stroke and dementia

Cross-trait genetic correlation between the two sub-classifications of WMH (deep and periventricular), stroke (overall and ischemic subtypes) and dementia were estimated using LDSC ^40^ on the GWAS summary statistics from phase II, MEGASTROKE (European ancestry-only)^41 42,43^. LD scores were based on the HapMap3 European reference panel. The significance threshold was set to 0.0026 (0.05/19 tests).

Regional level correlation was investigated using the GWAS-pairwise (GWAS-PW) method ^44^. GWAS-PW is a Bayesian approach that estimates the probability of a shared locus harboring genetic variant that influences only the first trait (model 1), only the second trait (model 2) or both traits (model 3). The posterior probability of association (PPA) for model 3 greater than 0.90 was considered significant. Additionally, fgwas v.0.3.6^45^ with the default settings was run to determine the PPA for each of the traits separately and the overall correlation in Z-scores estimated from genomic regions with PPA less than 0.20 was used to correct for possible sample overlap between the traits.

## Results

### Demographic characteristics

Study descriptions are provided in the Supplementary Information and Supplementary Table 1. The discovery cohort was comprised of dementia and stroke free older adults (≥45 years, n ∼18,234) from 16 international studies. The majority of the sample had DWMH and PVWMH data extracted by automated or semi-automated volumetric methods (n = 14,383), while the remainder used visual ratings (Supplementary Table 2). The discovery cohort was primarily Caucasian but included a smaller subsample of African Americans (n=736) and Hispanics (n= 658). The replication cohort consisted of predominantly Caucasian participants from the UK Biobank (n=8,428), all of whom had DWMH and PVWMH extracted using automated methods. Genotyping, imputation and analysis details for each study are provided in Supplementary Table 3.

### Discovery and Replication

The discovery association meta-analysis (Phase I) found genome-wide significant associations in the 17q25.1 region for both DWMH (n=167 SNPs) and PVWMH (n=199 SNPs; Supplementary Tables 4 and 5). This region has previously been associated with total WMH burden^15,16,19^. In contrast, only the PVWMH analysis found additional genome-wide significant associations on chromosomes 2 (3 *NBEAL* intronic SNPs) and 10 (2 regions, [i] a previously undescribed locus at 10q23.1 -14 *TSPAN14/FAM231A* intronic & intergenic SNPs; [ii] locus at 10q24.33 - one *SH3PXD2A* intronic SNP). Two of these regions have also previously been described for total WMH burden (chr 2, *NBEAL*^16,19^, 10q24.33, *SH3PXD2A*^19^). We repeated our analysis adjusting for hypertension, a known risk factor for WMH and cerebral SVD; this adjustment made almost no difference to our findings (Supplementary Tables 8 and 9). Replication of the majority of genome-wide significant results using participants from the UKB for both DWMH and PVWMH was observed after Bonferroni adjustment for multiple testing (DWMH *p* <3.6e-4, PVWMH *p* < 2.76e-4, Supplementary Tables 6 and 7).

### Overall Meta-analysis

Given the relatively large size of the replication cohort, a combined meta-analysis (Phase II) was undertaken using all of the available samples from the CHARGE, ENIGMA and UK Biobank cohorts (n∼26,654). Removing either the small subsample of non-Caucasians, or the cohorts with visual ratings, did not substantially change the findings for either phenotype (beta value r^2^ > 0.93).

Results of the Phase II GWAS meta-analysis for DWMH and PVWMH are shown in Table 1, Figure 1 and Supplementary Tables 10 and 11. We identified 513 genome-wide significant SNPs for PVWMH, and 236 for DWMH. In total, 209 SNPs were found to be significant for both phenotypes, all of which are located in 17q25.1 (Figure 2a).

**Figure 1.**
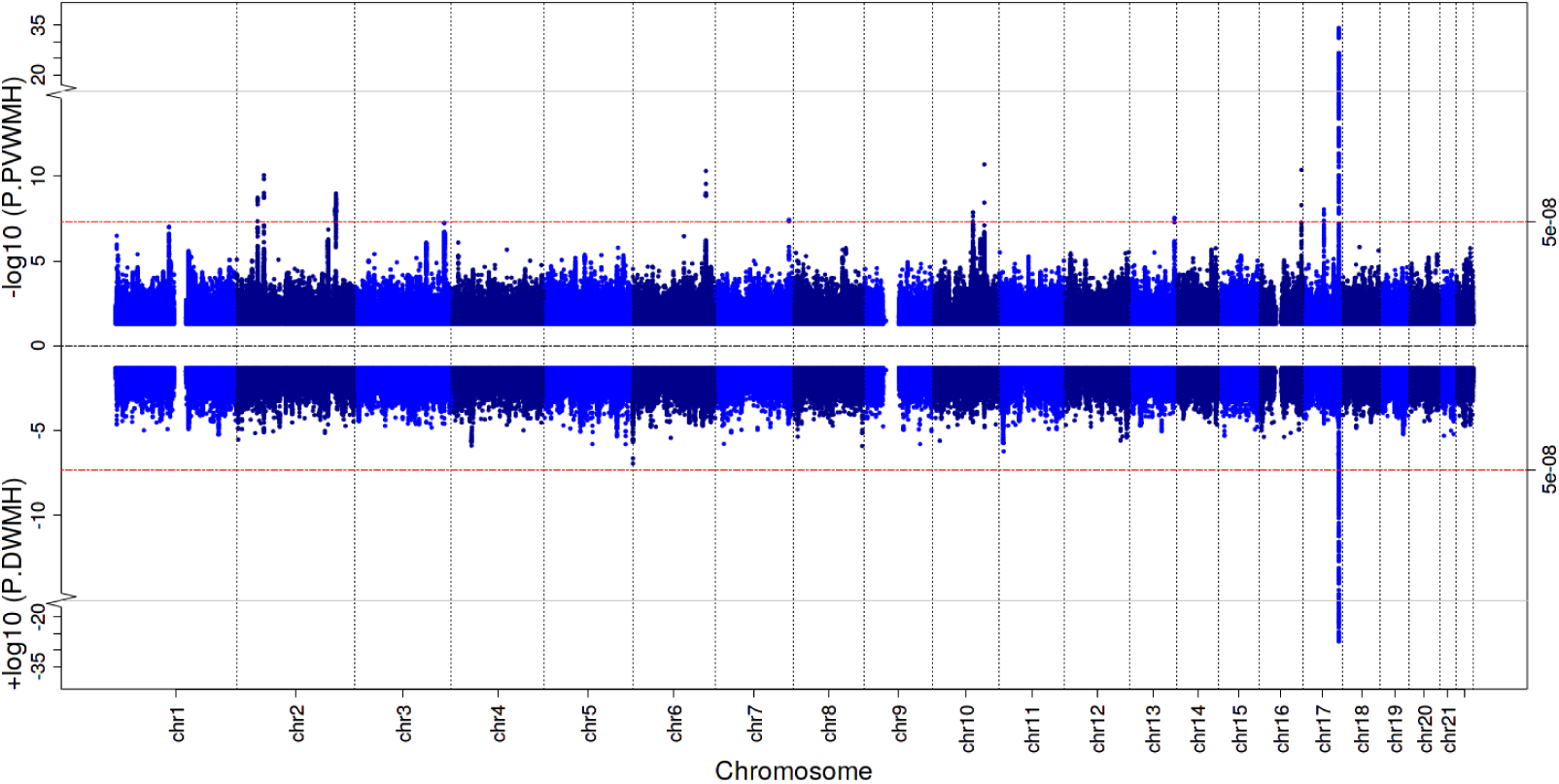
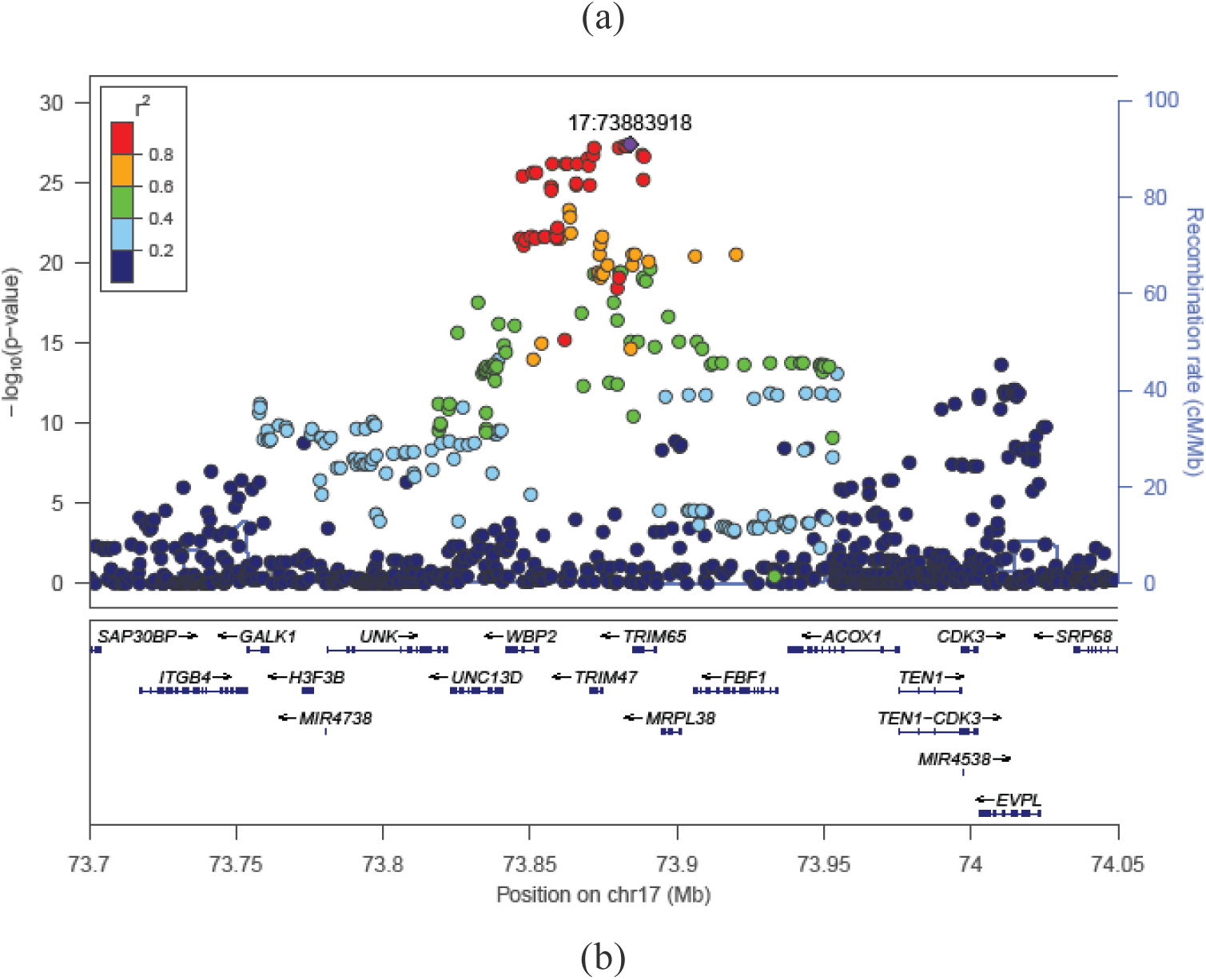
(a) Phase II GWAS meta-analysis results (n=26,654). Miami plot for periventricular WMH (PVWMH, upper panel) and deep WMH (DWMH, lower panel). Red dashed line shows genome-wide significance threshold (*p*<5e-8). (1b) Chr17 regional plot of genome-wide significant SNPs for DWMH. Colors of the SNPs indicate the level of LD with the top SNP (purple), rs35392904.

**Figure 2.**
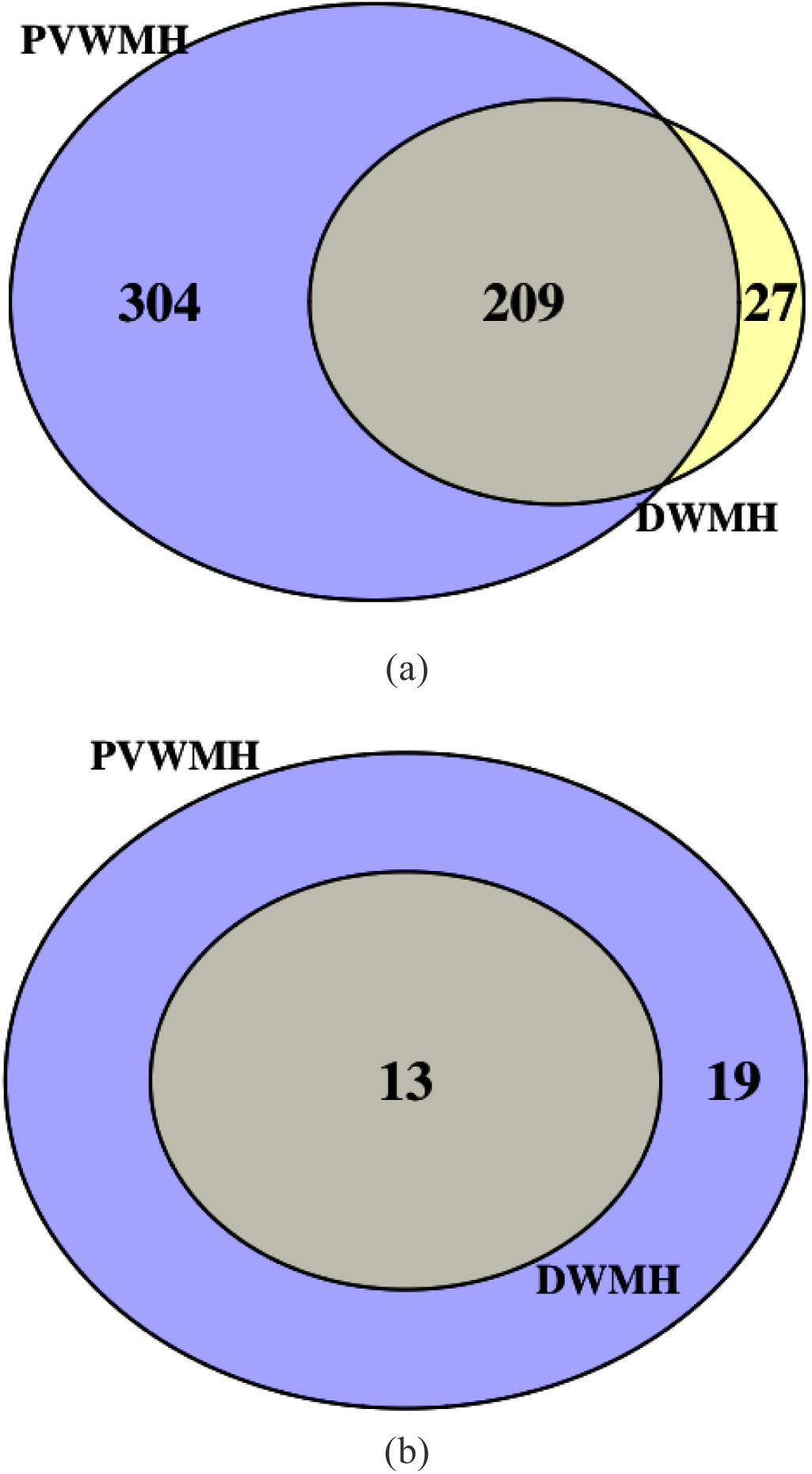
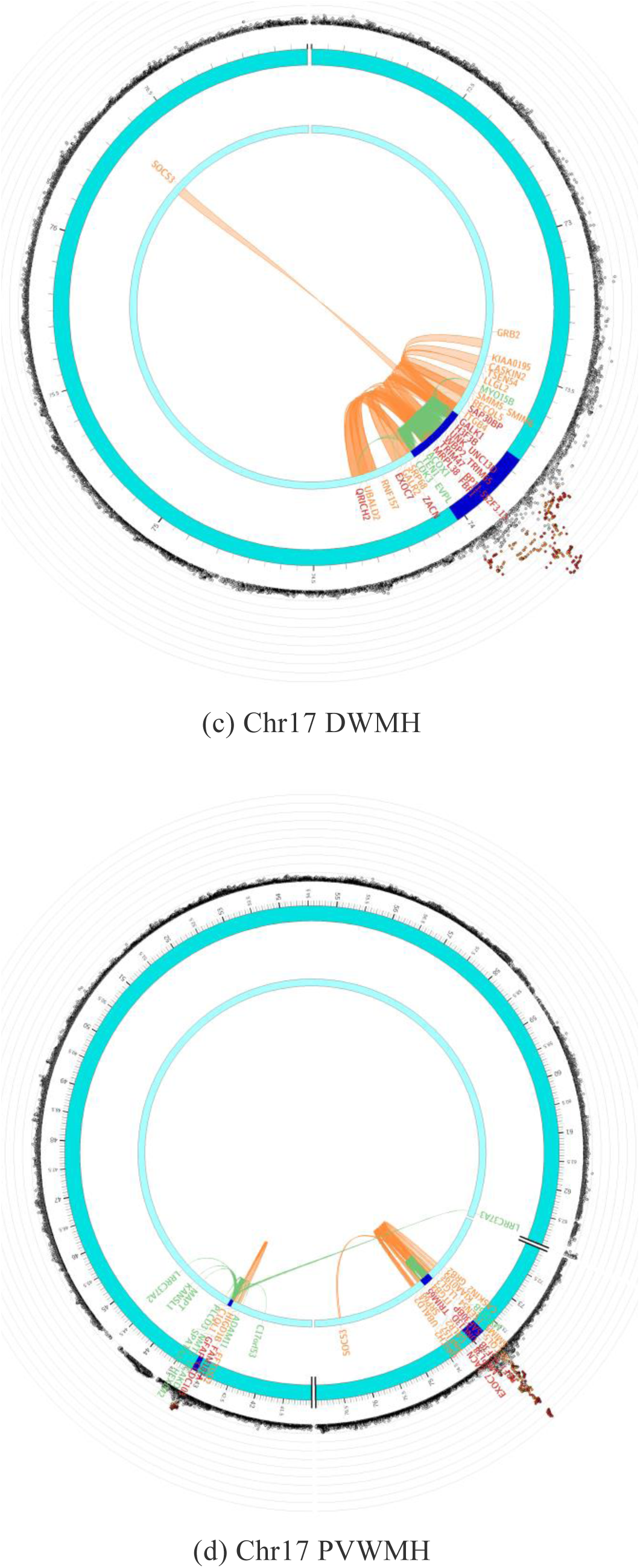
(a) Overlap between genome-wide significant SNPs (*p*<5e-8) for DWMH and PVWMH. (b) Overlap between significant genes identified by MAGMA for both phenotypes. (c - d) Circos plots for chr17 for both phenotypes, showing one identified region for DWMH but two for PVWMH. Outer ring shows SNPs <.05 with the most significant SNPs located towards the outermost ring. SNPs in high LD with the independent significant SNPs in each locus are colored in red (r^2^>.8)-blue (r^2^>.2); no LD (grey). Genomic risk loci are colored in dark blue (2^nd^ layer). Genes are mapped by chromatin interaction (orange), eQTL (green) or both (red).

**Table 1.**
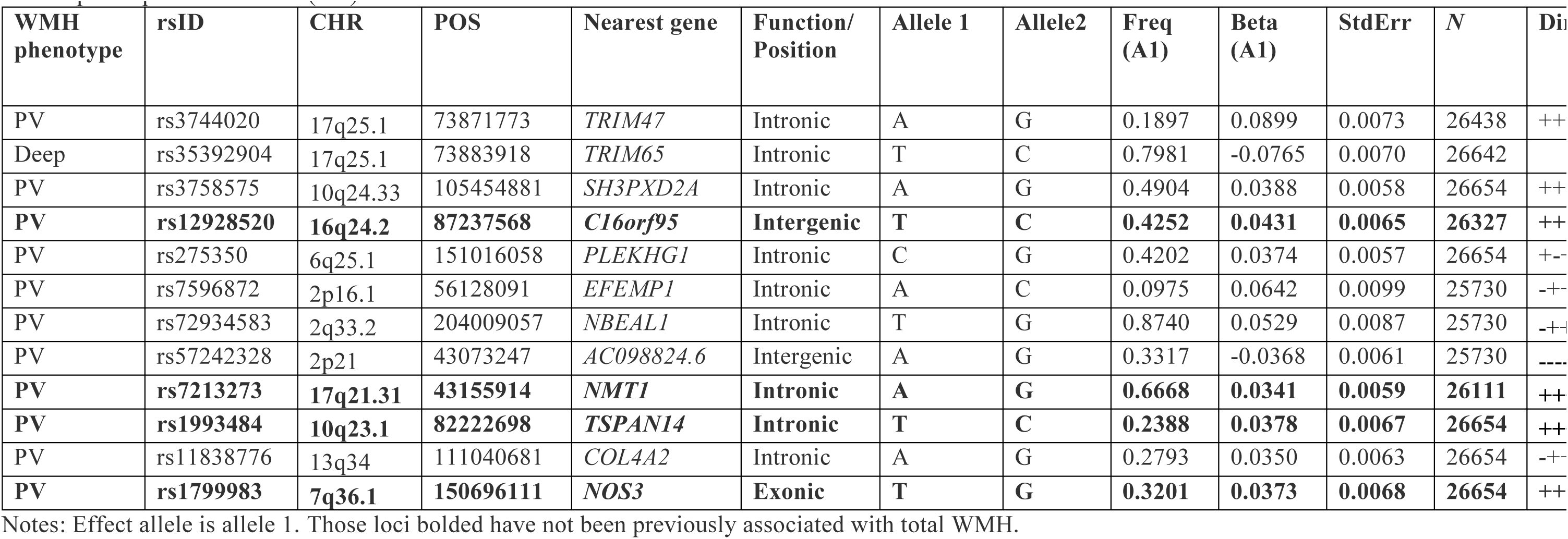
Top genome-wide significant SNP results from each genomic locus identified from the Phase II GWAS meta-analysis for deep and periventricular (PV) WMH.

### New genome-wide significant PVWMH loci

Ten chromosomal regions containing 290 genome-wide significant SNPs for PVWMH only were identified on chromosomes 2 (3 regions), 6, 7, 10 (2 regions), 13, 16 and 17q23.1 (Supplementary Table 11; Supplementary Figure 1). Four loci had not been previously reported for associations with total WMH at the genome-wide significant level. These new loci included a novel region (7.2kb) containing 2 genome-wide significant exonic SNPs in the *NOS3* gene on 7q36.1, with the top SNP (rs1799983) resulting in an amino-acid change, which is predicted to be tolerated. A new region on chr 10 was identified (10q23.1, 50.5kb, 4 intronic SNPs in *TSPAN14* & *FAM231A*) as well as one on 16q24.2 (1.2kb) which contained two genome-wide significant intergenic SNPs. An additional novel region to the 17q25.1 locus was also observed for PVWMH only. This 27.2kb region at 17q21.31 contained 8 genome-wide significant SNPs, most of which were intronic and found in the *NMT1* gene. Many of these are eQTLs or participate in chromatin interactions (Figure 2d).

Regions previously linked to total WMH burden were significant only for PVWMH in our analysis, which included three separate genomic loci on chromosome 2. The first region (2p16.1) spanned 27.3 kb and contained 6 genome-wide significant SNPs, with the top SNP (rs7596872, intronic, *EFEMP1*) having been reported previously^29^. The second region (2q33.2) spanned 442.3kb containing 250 significant SNPs, with the top SNP being rs72934583, located in an intron of *NBEAL1*. This SNP is an eQTL for several genes (*CARF, ICA1L, NBEAL1, FAM117B*) in various tissues. The third region (2p21) contained 7 SNPs, spanning 7.2 kb, with the top SNP, rs57242328, being intergenic and previously reported as an eQTL for *OXER1* (tibial nerve). Many SNPs in this region were suggestive (p<1e-5, n=215, Supplementary Table 11). Of the genome-wide SNPs, twelve were located in two regions (2p16.1, 2q33.2) previously associated with total WMH burden (rs78857879, rs7596872 – both *EFEMP1* intronic; rs72934505- *NBEAL* intronic), coronary heart disease, myocardial infarction, migraine and cholesterol (Supplementary Table 13). The third locus at 2p21 has also previously been significantly associated with total WMH burden (rs11679640), although this SNP was in low LD (r^2^=0.232) with our top SNP in this region (rs57242328)^19^.

The region on 6q25.1 contained six genome-wide significant SNPs spanning a region of 7.7 kb. The top SNP, rs275350, is an intronic SNP in *PLEKHG1*, which has been reported as an eQTL for *PLEKHG1* (testis) and recently associated with total WMH^29^. A region on 10q24.33 (4.3kb, 2 intronic SNPs in *SH3PXD2A*) was previously reported by Verhaaren et al^19^, who observed 4 genome-wide significant intronic SNPs in the *NEURL, PDCD11* and *SH3PXD2A* genes. Additionally, two genome-wide SNPs were located in the *COL4A2* gene (intronic) on 13q34 (spanning 2.6kb); all of these SNPs are in high LD (R^2^>0.8). In prior GWAS these SNPs have been associated with coronary artery disease (rs11838776) and myocardial infarction (rs55940034) (Supplementary Table 13).

### Genome-wide significant locus for DWMH and PVWMH

As expected, the association of the 17q25.1 locus with both phenotypes was confirmed. The size of this region including genome-wide significant SNPs only was similar for both DWMH (236 SNPs, BP 73757836-74025656, Figure 1b) and PVWMH (223 SNPs, BP 73757836-74024711, Supplementary Figure 1a). The top genome-wide significant results in this locus were rs3744020 for DWMH, (*p*=7.06e-35, intronic SNP, *TRIM47*) and rs35392904 for PVWMH (*p*=3.989e-28, intronic SNP, *TRIM65*), which are in high linkage disequilibrium (LD, R^2^=0.902) (Table 1). The top SNP for DWMH, rs3744020 was also reported genome-wide significant in the most recent total WMH GWAS^29^.Both SNPs (rs3744020, rs35392904) have been reported as eQTLs for several genes, including *TRIM47* and *TRIM65* in a range of tissues, including the cerebellum and cortex (GTEx^36^). Of the genome-wide significant SNPs, eight have previously been identified in GWAS for total WMH burden, psoriasis, systolic blood pressure or eosinophil and basophil cell counts (Supplementary Tables 12 and 13). Many of these SNPs are eQTLs or have chromatin interactions (Figure 2c-d).

### Gene-based association analysis

Using FUMA^33^, 13 genes reached genome-wide significance (*p*<2.66e-6) for association with both DWMH and PVWMH (Table 2, Supplementary Tables 14-15, Figure 2b). All of these genes are located at 17q25.1. For PVWMH, an additional 19 genes (Table 2, Supplementary Table 15) reached genome-wide significance, covering the majority of regions/loci found in the SNP-based analysis (apart from 2p16.1 and 2p21). Four genes were located in previously unidentified regions: *CALCRL* (2q32.1), *KLHL24* (3q27.1), *VCAN* (5q27.1) and *POLR2F* (22q13.1).

**Table 2.**
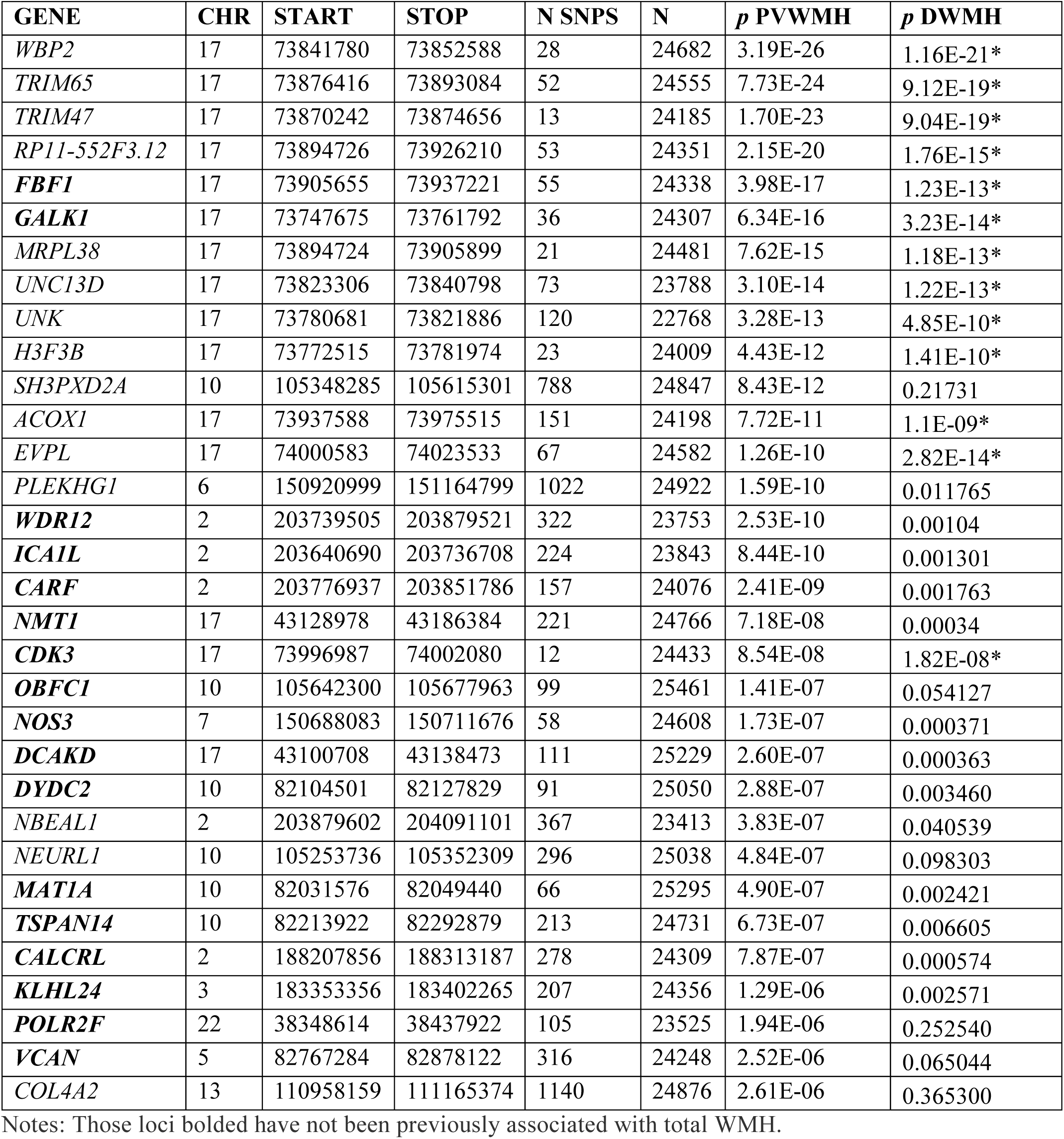
Thirty-two significant genes were identified for PVWMH using gene-based tests (*p*<2.66e-6). Thirteen of these genes (chr17) were also significant for DWMH (*).

### Pathway Analysis

We tested enrichment of MAGMA-derived gene-based *p*-values from the phase II analysis, in gene sets from MsigDBv6.1, as implemented in FUMA^33^. No significant associations with PVWMH were found, and only one gene set (*caffarel_response_to_thc_8hr_5_dn*) remained significant after multiple testing correction for DWMH.

### Heritability

Applying LD score regression^40^ to the phase II GWAS summary statistics, heritability was estimated to be 0.1 for DWMH and 0.18 for PVWMH. For the Rotterdam study, the largest individual cohort in Phase I, we used GCTA^39^ on the individual level data, adjusting for age and sex. We found heritability estimates of 0.29 and 0.47 for DWMH and PVWMH respectively.

### Genetic correlation with other traits

The genetic correlation between DWMH and PVWMH, estimated using the phase II GWAS results, was high (r_g_= 0.927), indicating a shared genetic architecture (Supplementary Table 16). Since higher WMH burden is associated with the presence of stroke and dementia, we also examined their genetic correlation with DWMH and PVWMH (Figure 3). Positive genetic correlations with both phenotypes were found for ‘all stroke’, ischemic stroke and SVD. Intracerebral haemorrhage (ICH, all types) was correlated with DWMH only. Deep ICH was correlated with both DWMH and PVWMH, but the relationship was much stronger for DWMH. No significant correlation was found with dementia.

**Figure 3.**
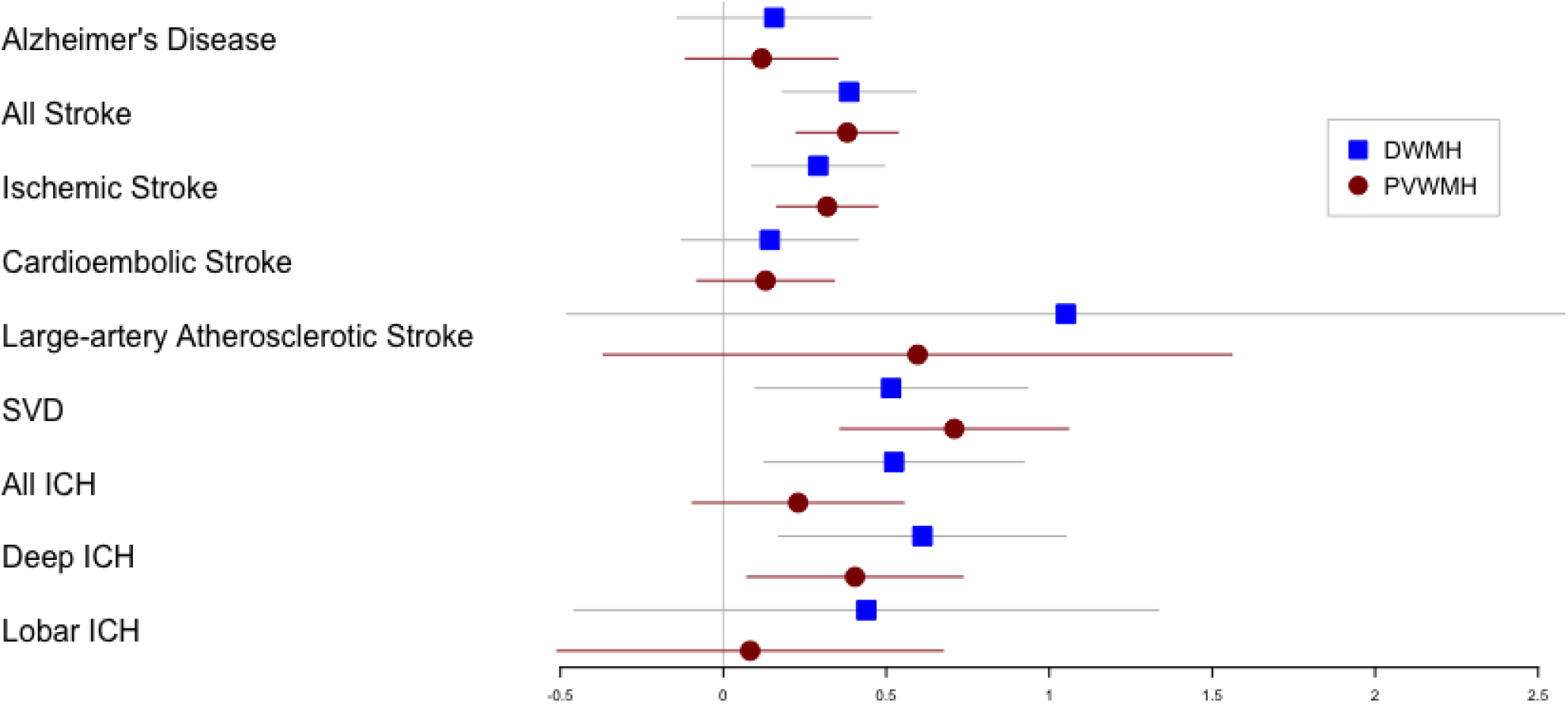
Genetic correlations (r_g_) between deep WMH (DWMH), periventricular WMH (PVWMH), Alzheimer’s disease (AD) and stroke phenotypes. Horizontal bars represent standard errors and the size of the square corresponds precision. SVD = small vessel disease stroke, All ICH = All intracranial hemorrhage, Deep ICH = deep intracranial hemorrhage, Lobar ICH = lobar intracranial hemorrhage

Using GWAS-PW^46^ we investigated if there were any genomic regions with shared genetic influence between PVWMH and DWMH. We identified several regions, including the major locus on chromosome 17 as well as chromosome 2 (two regions), 6 and 16 (Supplementary Table 17). These regions of shared influence overlap several genome-wide significant loci that were identified for PVWMH. However, using an alternative method^47^ to calculate local shared influence, only one region was significant, and was located within the 17q25.1 region (p=4.6e-6) (Supplementary Table 17).

Finally, we investigated local regions of shared influence between the WMH subtypes and stroke (Supplementary Table 17). A region on chromosome 7 (BP 149861853-150709571, *GIMAP4/KCNH2*) exhibited shared genetic influence of ‘all stroke’ with both DWMH and PVWMH. This region encompasses the PVWMH genome-wide significant *NOS3* exonic SNP. Other regions of shared influence with all stroke were observed for PVWMH only. For the sub-types of stroke (i.e. ischemic, cardioembolic and SVD), significant regions were identified for DWMH and PVWMH, but none were found for both phenotypes except the chromosome 7 region for ischemic stroke (also identified for all stroke). Additional regions of shared genetic influence were observed for PVWMH with ischemic stroke, cardioembolic and SVD. In contrast, DWMH shared a region of genetic influence with SVD stroke only (chr 6: BP 202452-1452004, *FOXF2/FOXQ1*). Using HESS^47^, no regions were found to have significant correlation between DWMH and any of the stroke subtypes. However, PVWMH had suggestive regions of genetic correlation with cardioembolic stroke (chr7, unadjusted p=0.03) and SVD (chr13, unadjusted p=0.02, *COL4A1/COL4A2*; chr2, unadjusted p=0.02, *NBEAL1*).

## Discussion

In our analysis using all available individuals (n=26,654, Phase II analysis), we found common and different results for DWMH and PVWMH burden. Newly discovered significant genes for PVWMH include *CALCRL, VCAN* and *NOS3*. The only significant locus (17q25.1) for DWMH was also found for PVWMH, which was the previously reported major locus for total WMH burden^15,16,19^. We found significant associations with PVWMH for all previously reported loci associated for total WMH. Our meta-analysis discovered a high number of significant SNPs, genes and loci that are only associated with PVWMH, more than any single total WMH GWAS to date. Most genes and loci previously reported as significant in total WMH^29,30,32,37,39^ were found to be associated with PVWMH alone, including *PLEKHG1*^*29*^, *SH3PXD2A*^*17,41,48*^ and *COL4A2*^*48*^. Most of the genes associated with PVWMH have direct effects on vascular function, or with vascular diseases such as ischemic stroke, intracranial hemorrhage or coronary artery disease. Other genes discussed as potential candidates in prior total WMH studies we now find to be significantly associated only with PVWMH including *DYDC2* and *NEURL1* as well as *NMT1, GALK1, H3F3B, UNK, UNC13D, EVPL, ICAL1, WDR12/CARF, NBEAL1*, and *EFEMP1*.

Genetic correlation analyses between DWMH and PVWMH and other phenotypes indicate independent and shared genetic architecture. The genetic correlation between the two phenotypes was high, as previously reported in a twin analysis^13^. Regional overlap analyses demonstrated shared genomic regions between PVWMH and DWMH at 2p16.1 (*EFEMP1*), 2q33.2 (*CARF* & *NBEAL*), 6q25.1 (*PLEKHGI*^*29*^), 16q24.2 (*C16orf95*), and 17q25.1 (*TRIM47, TRIM65*). Comparison of these phenotypes with ischemic and hemorrhagic stroke phenotypes from the MEGASTROKE Consortium^41^ for shared genetic architecture yielded interesting results. In general, DWMH and PVWMH both had shared and unique associations with different genomic regions and stroke phenotypes, including “any stroke” and “ischemic stroke”. For the first time, we demonstrated shared genetic architecture between any stroke, SVD stroke and PVWMH, at loci on chr 2 (*NBEAL1*), 3 (*PLSCR1*) and 10 (*SORBS1*). Significantly we discovered overlap for PVWMH only with deep intracranial hemorrhage (ICH) at genomic regions containing the genes *COL4A1/COL4A2*^49^ (chr13) and *CHD6* (chr20). There was also overlap for PVWMH with SVD stroke at the chr 2 region, which includes the *NBEAL1* gene. Many of these genes are implicated in cerebral small vessel disease, such as *COL4A1/COL4A2* and *NBEAL1.* Deep ICH is a regional classification of ICH occurring in deep subcortical regions approximating the same brain region as PVWMH. Together, these results are suggestive of an underlying vascular pathophysiology influenced by variation in genes with influences on microvessels in the periventricular region that are associated with deep ICH, periventricular SVD, and periventricular ischemic stroke.

We identified 11 independent loci for PVWMH, and one locus for DWMH. The only significant locus observed for DWMH was the previously reported total WMH 17q25.1 locus^15,16,19,29^, which was also found for PVWMH. Several of the identified loci overlap with prior total WMH GWAS results, specifically 17q25.1, 2p16.1, 2p21, 2q33.2, 6q25.1, 10q24.33 and 13q34^15,17,19,29^. Many of the previously reported candidates (17q25.1) achieved genome-wide significance for the first time in both PVWMH and DWMH including *TRIM47, TRIM65, MRPL38, WBP2, FBF1*, and *ACOX1*^*15-17,19*^ indicating shared biology. For both phenotypes, the majority of the top SNPs from each locus were intronic. As discussed, several of these SNPs are reported cis eQTLs in various tissues. There were many genes located within 10kb of our significant SNPs (Supplementary Tables 9 and 10). Fourteen of these genes overlapped with the two phenotypes and are located in the gene-rich locus, 17q25.1, including *EVPL, UNK*, and *CDK3*. Interestingly, in a post-mortem study comparing gene expression in DWMH versus non-lesion control white matter, *EVPL* was downregulated in DWMH^50^. Pathway analyses did not find any significant results for PVWMH and only one pathway for DWMH was identified, which was not informative with respect to WMH.

PVWMH had significant associations with loci containing genes implicated in large and small vessel disease, as well as ischemic and hemorrhagic stroke. The association of these vascular genes with PVWMH and not DWMH suggests a potential underlying regional difference in the microvasculature affecting the risk of vascular injury in the periventricular region. The different physiology of the short penetrating microvessels emerging from the midline arteries of the brain unique to the periventricular region may provide explanations for our observed differences when comparing the two phenotype results ^1,7,21,23^. Some of these genes such as *SH3PXD2A* have been previously associated with total WMH and ischemic stroke^16,41^. The *NOS3* gene is associated with coronary artery disease, migraine, vascular dysfunction, SVD, and ischemic stroke^29,43,51,52^. *PLEKHG1* is associated with dementia and ischemic stroke^17,53^. The most notable associated vascular gene is *COL4A2* that encodes for a subunit of type IV collagen, which has been associated with SVD, ischemic stroke, intracranial hemorrhage, and coronary artery disease^17,49,54-56^. It is a proposed therapeutic target for the prevention of intracranial hemorrhage^57,58^. Verhaaren et al.^19^ identified a suggestive intronic *COL4A2* SNP for total WMH, rs9515201, which was genome-wide significant in Traylor et al.^29^. Our genome-wide significant top SNP (intronic) in the *COL4A2* gene, rs11838776, is in high LD with rs9515201 (r^2^=0.83). New vascular genes not previously reported associated with total WMH, include the *MAT1A* gene, which catalyzes adenosine transfer from ATP to methionine and is associated with stroke and hypertension^59^. *FAM213A* encodes for the peroxiredoxin like 2A protein, which is an antioxidant associated with endothelial activation^60^. The *CALCRL* gene encodes the calcitonin receptor like receptor and modulates the calcitonin signaling system, implicated in ischemic stroke ^61^.

Other newly significant non-vascular PVWMH genes identified by gene-based analysis include *TSPAN14* (10q23.1), which encodes one of the tetraspanins, which organize a network of interactions referred to as the tetraspanin web. This includes ADAM10 - a metalloprotease that cleaves the precursor of cell surface proteins, including the amyloid precursor protein and Notch^62^. Another newly identified gene, *VCAN* (5q14.2-q14.3), encodes a large chondroitin sulfate proteoglycan that is found in the extracellular matrix. In a recent meta-analysis, *VCAN* was associated with white matter microstructural integrity in UK Biobank participants with WMH, stroke, and dementia^63^. *KLHL24* (3q27.1), encodes a ubiquitin ligase substrate receptor that is regulated by auto-ubiquitination and is associated with deficits in skin keratization^64^. These candidate genes for PVWMH may influence the immediate tissues surrounding microvessels and may contribute to SVD-associated biological changes.

DWMH and PVWMH share a previously reported locus on 17q25.1 for total WMH, with these SNP associations having the largest effect sizes for both phenotypes. Thus, this locus appears to influence both WMH subtypes and is the most predominant genetic association with the largest potential clinical effect for both phenotypes. The top genome-wide significant hits for DWMH and PVWMH (17q25.1) were either identical with the SNP recently reported by Traylor et al^29^for total WMH (PVWMH SNP rs3744020) or in high LD (R^2^>0.9) with the previously identified top ranked SNPs from prior total WMH GWAS in the same locus (rs3744028, Fornage et al.^15^, rs7214628, Verhaaren et al.^19^). These SNPs were only in moderate LD (R^2^≤0.396) with the top SNP (rs3760128) identified in a recent exome association analysis ^16^. All of these SNPs fall within or between the previously reported *TRIM47* and *TRIM65* genes^15,17,19,29^. This gene-rich locus contains genes that influence glial cell proliferation and have been hypothesized to influence gliosis, which is a histological and MRI marker of microvascular injury^1^. It includes previously identified total WMH genes, such as *TRIM47/TRIM65* (glial proliferation, astrocytoma’s)^15,19^, the *ACOX1* gene (cell replication, hepatic cancer)^15,16,19,65^ and the *MRPL38* gene (protein synthesis)^16^. Genes associated with neuronal injury and/or neurodegenerative disorders are also found in the 17q25.1 locus, including *CDK3* (neuronal cell death in stroke)^66^, *H3F3B* (schizophrenia pathogenesis)^67^ and the *GALK1* gene (galactosemia, a neurodegenerative metabolic disease)^68^. Interestingly, two genome-wide significant intronic *UNC13D* SNPs identified in this study and reported previously for total WMH burden^19^, rs9894244 and rs7216615, have been reported as eQTLs for *GALK1* and *H3F3B* respectively^*69*^.

The PVWMH specific loci also contained genes that potentially influence astrocytic function and gliosis. There were several previously reported total WMH candidate genes such as *NBEAL1* (2q33.2, astrocytoma)^16,19^, *WDR12* (2q33.2, cell cycle, apoptosis, myocardial infarction, coronary artery disease, carotid intima-media thickness)^16^, *NEURL1* (10q24.33, notch signaling, cell proliferation in medulloblastoma, atrial fibrillation, total WMH^15,16,19^), *CARF* (2q33.2, modulates BDNF expression, tumor suppression, astrocyte function^70^) and *EFEMP1* (2p16.1, malignant gliomas^54^). Newly identified genes include the *NMT1* gene (17q21.31), which catalyzes the transfer of myristate from CoA to proteins^71^. The *ICA1L*, islet cell autoantigen 1-like protein gene, approximating CARF, is associated with amyotrophic lateral sclerosis and may influence neuronal death in that disease^72^. *POLR2F* (22q13.1) is associated with colon cancer^73^ while *OBFC1* (10q24.33) initiates DNA replication^74^ and *DYDC2* (10q23.1) is associated with myofibrillar myopathy and ventricular cardiomegaly^75^.

While our GWAS were only slightly larger than the previous biggest GWAS on total WMH burden with 21,079 participants^19^, our detection rate of significant SNPs was substantially higher. This improved detection may be the result of reduced heterogeneity by separately analyzing the DWMH and PVWMH phenotypes. Additionally, sensitivity also may have been increased by including more modern samples (e.g., from cohorts such as UK Biobank), with most data acquired on the more powerful 1.5-3.0 Tesla MRI scanners using sequences such as FLAIR and incorporating segmentation programs with better lesion detection and volumetric quantification. The results obtained did not change significantly when cohorts with visual rather than volumetric measures were excluded from the analysis.

Shortcomings of this study include the potential presence of phenotype heterogeneity due to the different WMH extraction algorithms used, with a minority of samples using visual ratings. However, this is a common problem encountered in this type of study^15,16,19^. The majority of the participants in this study were Caucasian and hence these results may not apply to other ethnicities. Sex differences have been previously reported but were also not examined^76^. For the Phase II meta-analysis, we did not have an independent replication cohort. Even though our results suggest improved power and reduction in potential bias through the discrimination of PVWMH from DWMH, the Euclidean methodology used by the majority of studies, which prescribed PVWMH as all WMH within one centimeter of the ventricle walls, undoubtedly missed PVWMH lesions outside this boundary. It is also possible that a number of DWMH lesions were accidentally included although review of the data makes this seem less likely. There are unique “cuff regions” or “border zone regions”, which fall within the PVWMH phenotype that represent a lesion type that is specific to the regions abutting the anterior and posterior horns of the lateral ventricles. These lesions have been attributed with an as yet not fully classified pathophysiology that could be quite different from PVWMH and DWMH. This could represent a hitherto unclassified source of phenotype heterogeneity unaccounted for in the present analysis. Further, we did not investigate other WMH sub-classifications or alternative measures. For example, other regional sub-classifications may prove fruitful (e.g. lobar).

## Conclusion

When using the two anatomically based sub-classifications (PVWMH and DWMH), we identified new genetic associations with WMH. Novel genetic associations unique to PVWMH alone were found and include genes such as *CALCRL, KLHL24, VCAN, MAT1A, OBFC1, DCAKD*, and *POLRF2*. We observed that the PVWMH phenotype was associated with genes previously associated with vascular phenotypes such as SVD, ICH and ischemic stroke. We found shared genetic architecture between PVWMH and deep ICH that includes loci containing the genes *COL4A1/COL4A2* and *CHD6*, supporting a strong etiologic association between these clinical phenotypes and the periventricular region. Our findings identify underlying cellular processes that are involved in the response to chronic brain injury. These include genes implicated in astrocytic/glial and vascular responses. Finally, we discovered a new set of putative PVWMH genes including *TSPAN14* and *VCAN*, which suggest new mechanisms that may influence the connective tissue outside the microvasculature. In conclusion, our findings suggest that the two WMH subclassifications have shared (17q25.1) but also unique and divergent genetic underpinnings. This work provides stimulus for further study to assess the roles of the identified genetic variants and genes in diseases of the aging brain.

