## Supplementary Notes for "Common genetic variation indicates separate etiologies for periventricular and deep white matter hyperintensities"

**Supplementary File**

1. Cohort descriptions
2. Cohort Acknowledgements and Funding
3. Supplementary Figures

*Supplementary Figure 1*. Regional Plots of genome-wide significant SNPs for periventricular WMHs

*Supplementary Figure 2.* Circos plots for each chromosome containing significant loci.

4. Supplementary Table Legends
1. **Cohort Descriptions**

*Age Gene-Environment Susceptibility - Reykjavik Study* (AGES-Reykjavik)

The AGES-Reykjavik Study is a single center prospective cohort study based on the Reykjavik Study. The Reykjavik Study was initiated in 1967 by the Icelandic Heart Association to study cardiovascular disease and risk factors. The cohort included men and women born between 1907 and 1935 who lived in Reykjavik at the 1967 baseline examination. Re-examination of surviving members of the cohort was initiated in 2002 as part of the AGES-Reykjavik Study. The AGES-Reykjavik Study is designed to investigate aging using a multifaceted comprehensive approach that includes detailed measures of brain function and structure. All cohort members were European Caucasians. The study design has been described previously^1^. Briefly, as part of a comprehensive examination, all participants answered a questionnaire, underwent a clinical examination and had blood drawn. All consenting participants without contraindications were offered a brain MRI on a dedicated machine in the study center: a total of 5003 participants had an MRI.2 Of these, 3664 were genotyped at the Laboratory of Neurogenetics, Intramural Research Program, NIA, Bethesda, Maryland, and 3219 participants passed QC criteria for genotyping. AGES–RS was approved by the National Bioethics Committee in Iceland (VSN 00-063) acting as the institutional review board for the Icelandic Heart Association, and by the institutional review board governing the National Institute on Aging. All participants gave written informed consent. More details on the design of the study can be found in Harris, et. al.^1^ .

*Atherosclerosis Risk in Communities Study (ARIC)*

The ARIC study is a population-based cohort study of atherosclerosis and clinical atherosclerotic diseases^2^. At its inception (1987-1989), 15,792 men and women, including 11,478 white and 4,266 black participants were recruited from four U.S. communities: Suburban Minneapolis, Minnesota; Washington County, Maryland; Forsyth County, North Carolina; and Jackson, Mississippi. In the first 3 communities, the sample reflects the demographic composition of the community. In Jackson, only black residents were enrolled. Participants were between age 45 and 64 years at their baseline examination in 1987-1989 when blood was drawn for DNA extraction and participants consented to genetic testing. Vascular risk factors and outcomes, including transient ischemic attack, stroke and dementia, were determined in a standard fashion. During the first 2 years (1993-1994) of the third ARIC examination (V3), participants aged 55 and older from the Forsyth County and Jackson sites were invited to undergo cranial MRI. This subgroup of individuals with MRI scanning represents a random sample of the full cohort because examination dates were allocated at baseline through randomly selected induction cycles.

*Austrian Stroke Prevention Family Study (ASPS-Fam)*

ASPS-Fam is a prospective single-center community-based study on the cerebral effects of vascular risk factors in the normal aged population of the city of Graz, Austria^3,4^. ASPS-Fam represents an extension of the Austrian Stroke Prevention Study (ASPS), which was established in 1991^5,6^. Between 2006 and 2013, study participants of the ASPS and their first-grade relatives were invited to enter ASPS-Fam. Inclusion criteria were no history of previous stroke or dementia and a normal neurologic examination. A total of 419 individuals from 176 families were included into the study. The number of members per family ranged from 2 to 6. The entire cohort underwent a thorough diagnostic workup including clinical history, laboratory evaluation, cognitive testing, and an extended vascular risk factor assessment. They were all European Caucasians. Those 278 participants who passed genotyping quality control and underwent MRI scanning were available for these analyses.

*Erasmus Rucphen Family study (ERF)*

The Erasmus Rucphen Family study (ERF) is a family based cohort study that includes inhabitants of a genetically isolated community in the South-West of the Netherlands, studied as part of the Genetic Research in Isolated Population (GRIP) program^7,8^. The goal of the study is to identify the risk factors in the development of complex disorders. The study population includes approximately 3,000 individuals who are living descendants of 22 couples who lived in the isolate between 1850 and 1900 and had at least six children baptized in the community church. All data were collected between 2002 and 2005. All participants gave informed consent, and the Medical Ethics Committee of the Erasmus University Medical Centre approved the study.

*Framingham Heart Study (FHS)*

The FHS is a three-generation, single-site, community-based, ongoing cohort study that was initiated in 1948 to investigate the risk factors for cardiovascular disease. The study includes three generations of participants: the Original cohort followed since 1948^9^; their Offspring and spouses of the Offspring, followed since 1971^10^; and children from the largest Offspring families enrolled in 2000^11^. The Original cohort enrolled 5,209 men and women who comprised two-thirds of the adult population then residing in Framingham, MA. Survivors continue to receive biennial examinations. The Offspring cohort comprises 5,124 persons (including 3,514 biological offspring) who have been examined approximately once every 4 years. The Third-generation cohort includes 4,095 participants with at least one parent in the Offspring Cohort. The first two generations were invited to undergo an initial brain MRI in 1999-2005, and for the Third-generation, brain MRI began in 2009. FHS participants had DNA extracted and provided consent for genotyping. All participants provided written informed consent, and the study was approved by the Boston University Medical Center Institutional Review Board.

*Genetic Study of Atherosclerosis Risk (GeneSTAR)*

GeneSTAR is a longitudinal cohort study designed to determine environmental, phenotypic, and genetic causes of premature cardiovascular disease^12,13^. Participants (n=3533) came from European- and African-American families (n=891) identified from 1983-2006 from probands with documented coronary disease prior to 60 years of age who were identified at the time of hospitalization in any of 10 Baltimore area hospitals. After providing written informed consent, apparently healthy siblings of the probands and offspring of the siblings and probands were screened for traditional coronary disease and stroke risk factors. A random subset of this cohort participated in a brain MRI study between 2009 and 2013. Siblings and offspring were excluded if they had: atrial fibrillation or symptomatic cardiovascular disease of any kind, history of chronic corticosteroid use, life-threatening diseases, neurologic diseases that would preclude accurate MRI interpretation, or implanted metals that prohibited MRI scans.

*HUNT*

The HUNT Study is a collaboration between the HUNT Research Centre (Faculty of Medicine and Movement Sciences, NTNU – Norwegian University of Science and Technology), Nord-Trøndelag County Council, Central Norway Health Authority, and the Norwegian Institute of Public Health.

*LIFE Adult*

LIFE Adult is a population-based study and a part of the large-scale research project LIFE (Leipzig Research Center for Civilization Diseases). 10,000 residents (main age range 40 – 79) from the district of Leipzig (Saxony, Germany) were recruited and extensively phenotyped for a number of disease and environmental parameters^14^. All subjects gave written informed consent to participate in the study. The procedures were conducted according to the Declaration of Helsinki and approved by the University of Leipzig's ethics committee (registration-number: 263-2009-14122009). Details of genotyping, imputation and corresponding quality control can be found elsewhere^15^.

*Lothian Birth Cohort 1936 (LBC1936), Scotland*

The LBC1936 consists of relatively healthy individuals assessed on cognitive and medical measures at age 70 years (n=1,091), and again with brain imaging traits at 73 years of age (n=866). They were born in 1936, most took part in the Scottish Mental Survey of 1947, and almost all lived independently in the Lothian region of Scotland. A full description of participant recruitment and testing can be found elsewhere^16-18^. The study was approved by the Lothian (REC 07/MRE00/58) and Scottish Multicentre (MREC/01/0/56) Research Ethics Committees and all subjects give written informed consent.

*Northern Manhattan Study (NOMAS)*

The Northern Manhattan Study (NOMAS) is a multi-ethnic population-based cohort study investigating risk factors for stroke and cognitive decline. A total of 3,298 stroke-free participants were enrolled in NOMAS between 1993 and 2001. Eligibility criteria included: 1) being clinically stroke-free, 2) aged >40 years old (>55 after 1998), and 3) living in Northern Manhattan for at least 3 months in a household with a telephone. From this original cohort, 1,091 participants were recruited for participation in the NOMAS Magnetic Resonance Imaging (MRI) Sub-Study. In addition, 199 household members were enrolled as previously described (Sub-Study Total *N*=1,290). The following eligibility criteria were used: a) >50 years old, b) no contraindications to MRI, and c) clinically stroke-free. All participants provided written informed consent and the study was approved by the Institutional Review Boards of Columbia University and the University of Miami. Further details of NOMAS can be found in Sacco et al, 2004^19^.

*Older Australian Twins Study (OATS)*

Older adult twins (65+ years) were recruited from Twins Registry Australia and through other avenues from the three eastern states of Australia into the Older Australian Twins Study (N=623). Exclusion criteria included a life-threatening illness or a current diagnosis of an acute psychiatric illness. A comprehensive assessment was undertaken including medical history, physical tests, demographics and blood was collected for genetic and biochemistry analyses. Neuroimaging was also undertaken on a subsample. Ethics approval was granted by Twins Registry Australia, University of New South Wales, University of Melbourne, Queensland Institute of Medical Research and the South Eastern Sydney and Illawarra Area Health Service, University of New South Wales and the Illawarra Area Health Service Human Research Ethics Committees. Written informed consent was provided by all participants. For further details, see Sachdev et al. 2009^20^.

*PROspective Study of Pravastatin in the Elderly at Risk (PROSPER)*

PROSPER was a prospective multicenter randomized placebo-controlled trial to assess whether treatment with pravastatin diminishes the risk of major vascular events in elderly. A detailed description of the study has been published elsewhere. Between December 1997 and May 1999, we screened and enrolled subjects in Scotland (Glasgow), Ireland (Cork), and the Netherlands (Leiden). Men and women aged 70-82 years were recruited if they had pre-existing vascular disease or increased risk of such disease because of smoking, hypertension, or diabetes. A total number of 5,804 subjects were randomly assigned to pravastatin or placebo. A large number of prospective tests were performed including Biobank tests and cognitive function measurements. Neuroimaging was performed in a random subset of 550 Dutch subjects. A whole genome wide screening has been performed in the sequential PHASE project. For 5,763 subjects, DNA was available for genotyping. The study was approved by the institutional ethics review boards of centres of Cork University (Ireland), Glasgow University (Scotland) and Leiden University Medical Center (the Netherlands) and all participants gave written informed consent^21-23^.

*Rotterdam Study (RS)*

The Rotterdam Study is a population-based cohort study among inhabitants of Ommoord, a district of Rotterdam, the Netherlands, and aims to examine the determinants of disease and health in the elderly^24^. In 1990-1993, 7,983 persons aged 55 years and older participated and were re-examined every 3 to 4 years (Rotterdam Study I). In 2000-2001 the cohort was expanded by 3,011 persons aged 55 and over who had not yet been part of the Rotterdam Study (Rotterdam Study II). In 2006-2008 a second expansion (Rotterdam Study III) of 3,932 persons aged 45 and over was realized. DNA was successfully extracted at baseline in 11,496 participants. In 1995 and 1999, MRI was performed in random subsamples from the Rotterdam Study, but was implemented in the core study protocol from 2005 onwards. More information regarding the MRI protocol is described elsewhere^25^. The Rotterdam Study has been approved by the Medical Ethics Committee of the Erasmus MC (registration number MEC 02.1015) and by the Dutch Ministry of Health, Welfare and Sport (Population Screening Act WBO, license number 1071272-159521-PG). The Rotterdam Study has been entered into the Netherlands National Trial Register (NTR; [www.trialregister.nl](http://www.trialregister.nl/)) and into the WHO International Clinical Trials Registry Platform (ICTRP; [www.who.int/ictrp/network/primary/en/](http://www.who.int/ictrp/network/primary/en/)) under shared catalogue number NTR6831. All participants provided written informed consent to participate in the study and to have their information obtained from treating physicians.

*Sydney Memory and Ageing Study (Sydney MAS)*

Participants aged 70-90 years of age were recruited randomly from the community in Sydney, Australia (N=1037) into a longitudinal study to investigate mild cognitive impairment and related syndromes and to determine the rate of cognitive change. Study exclusion criteria included diagnoses of dementia, schizophrenia or a progressive malignancy. An extensive interview was undertaken and questionnaire data collected, including demographics, cognitive performance and medical history. The majority of participants provided a blood sample for genetic and biochemistry analyses. Neuroimaging was performed on a subset of participants. The University of New South Wales and the Illawarra Area Health Service Human Research Ethics Committees gave ethics approval for the study. Written informed consent was provided by all participants. More information is provided in Sachdev et al. 2010^26^.

*Three-City Dijon (3C-Dijon)*

The 3C study is conducted in three French cities (Bordeaux, Dijon, and Montpellier), comprising 9,294 participants, designed to estimate the risk of dementia and cognitive impairment attributable to vascular factors. Eligibility criteria included living in the city and being registered on the electoral rolls in 1999, 65 years or older, and not institutionalized. The 3C-Dijon study recruited 4,931 individuals. The overall design of the 3C-Dijon study is detailed elsewhere. Participants aged less than 80 years and enrolled between June 1999 and September 2000 (n=2,763) were invited to undergo a brain MRI. Although 2,285 subjects agreed to participate (82.7%), because of financial limitations, only 1,924 MRI scans were performed. DNA samples of 3C-Dijon participants were genotyped at the Centre National de Génotypage, Evry, France, with Illumina Human610IQuad® BeadChips^27-29^.

*United Kingdom Biobank (UKB)*

The UKB is a large-scale longitudinal epidemiological study of over 500,000 individuals aged 40-69 years at baseline from the United Kingdom (http://www.ukbiobank.ac.uk). This richly phenotyped study includes the collection of blood samples for genetic analysis and brain imaging scans. The analyses presented here use data from the 2017 imaging data release of approximately 10K participants that were accessed via Data Application 8107. Genetic data are available for the majority of these individuals.

*Vietnam Era Twin Study of Aging (VETSA)*

Middle-age male twins in the VETSA were randomly recruited from the population-based Harvard Drug Study (HDS) between 2003-2007. WMH were assessed at VETSA wave 2. There were 1205 individual participants between the ages of 56 and 56 years at wave 2. WMH measures were available for 337 individuals. MRI began in year 3 of the original VETSA project, so those who entered the study earlier did not undergo imaging. MRI participants were demographically similar to the rest of the sample. Participants were concordant for US military service at some time between 1965 -1975. However, nearly 80% reported no combat experience. The sample is 88.3% non-Hispanic white, 5.3% African-American, 3.4% Hispanic, and 3.0% “other” participants. Based on data from the US National Center for Health Statistics, the sample is very similar to American men in their age range with respect to health and lifestyle characteristics. All analyses adjusted for correlated observations within twin pairs. The study was approved by Institutional Review Boards at the University of California, San Diego, Boston University, Massachusetts General Hospital, and the VA Puget Sound Healthcare System. Written informed consent was obtained from all participants.

2. **Funding and Acknowledgements**

*Age Gene-Environment Susceptibility - Reykjavik Study* (AGES-Reykjavik)

This study has been funded by NIH contracts N01-AG-1-2100 and 271201200022C, the NIA Intramural Research Program, Hjartavernd (the Icelandic Heart Association), and the Althingi (the Icelandic Parliament). The study is approved by the Icelandic National Bioethics Committee, VSN: 00-063. The researchers are indebted to the participants for their willingness to participate in the study.

*Atherosclerosis Risk in Communities Study (ARIC)*

The Atherosclerosis Risk in Communities Study is carried out as a collaborative study supported by National Heart, Lung, and Blood Institute contracts (HHSN268201100005C, HHSN268201100006C, HHSN268201100007C, HHSN268201100008C, HHSN268201100009C, HHSN268201100010C, HHSN268201100011C, and HHSN268201100012C), R01HL087641, R01HL59367 and R01HL086694; National Human Genome Research Institute contract U01HG004402; and National Institutes of Health contract HHSN268200625226C. The authors thank the staff and participants of the ARIC study for their important contributions. Infrastructure was partly supported by Grant Number UL1RR025005, a component of the National Institutes of Health and NIH Roadmap for Medical Research.

*Austrian Stroke Prevention Family Study (ASPS-Fam)*

The authors thank the staff and the participants for their valuable contributions. We thank Birgit Reinhart for her long-term administrative commitment, Elfi Hofer for the technical assistance at creating the DANN bank, Ing. Johann Semmler and Anita Harb for DNA sequencing and DANN analyses by TaqMan assays and Irmgard Poelzl for supervising the quality management processes after ISO9001 at the biobanking and DANN analyses. The research reported in this article was funded by the Austrian Science Fund (FWF) grant number P20545-P05, P13180 and PI904 as well as by the Austrian National Bank (OeNB) Anniversary Fund grant number P15435, the Austrian Federal Ministry of Science, Research and Economy under the aegis of the EU Joint Programme-Neurodegenerative Disease Research (JPND)-www.jpnd.eu and the Ph.D. program ‘Molecular Medicine’ of the Medical University of Graz. The Medical University of Graz supports the databank of the ASPS.

*Erasmus Rucphen Family study (ERF)*

The Erasmus Rucphen Family (ERF) study as a part of EUROSPAN (European Special Populations Research Network) was supported by European Commission FP6 STRP Grant No. 018947 (LSHG-CT-2006-01947) and also received funding from the European Community’s Seventh Framework Program (FP7/2007-2013)/Grant Agreement HEALTH-F4-2007-201413 by the European Commission under the program “Quality of Life and Management of the Living Resources” of 5th Framework Program (No. QLG2-CT-2002-01254). High-throughput analysis of the ERF data was supported by joint grant from Netherlands Organization for Scientific Research and the Russian Foundation for Basic Research (NWO-RFBR 047.017.043), and Russian Federal Agency of Scientific Organizations projects VI.53.2.2 and 0324-2015-0003. Exome sequencing analysis in ERF was supported by the Netherlands Organization for the Health Research and Development grant for Project No. 91111025. We thank all study participants and their relatives, general practitioners, and neurologists for their contributions and P. Veraart for help in genealogy, J. Vergeer for the supervision of the laboratory work, and P. Snijders for help in data collection.

*Framingham Heart Study (FHS)*

The Framingham Heart Study is supported by the National Heart, Lung and Blood Institute (NHLBI) Framingham Heart Study (contracts N01-HC-25195 and HHSN268201500001I), the Boston University School of Medicine, and by grants from the National Institute of Health (AG054076, AG033193, AG049505, AG033040, NS017950, AG049607, AG010129). We would like to thank the dedication of the Framingham Study participant. We also thank the Framingham study team, especially the Neurology core, both investigators and staff, for their contributions to data collection.

*Genetic Study of Atherosclerosis Risk (GeneSTAR)*

GeneSTAR was supported by grants from the National Institutes of Health National Institute of Neurological Disorders and Stroke (R01NS062059), the National Institutes of Health National Heart, Lung, and Blood Institute (U01 HL72518, HL087698), the National Institutes of Health/National Center for Research Resources (M01-RR000052) to the Johns Hopkins General Clinical Research Center, and National Institutes of Health/National Center for Research Resources and the National Center for Advancing Translational Sciences (UL1 RR 025005) to the Johns Hopkins Institute for Clinical and Translational Research. We would like to thank the participants and families of GeneSTAR and our dedicated staff for all their sacrifices.

*HUNT*

The HUNT Study is a collaboration between HUNT Research Centre (Faculty of Medicine and Movement Sciences, NTNU – Norwegian University of Science and Technology), Nord-Trøndelag County Council, Central Norway Health Authority, and the Norwegian Institute of Public Health. HUNT-MRI was funded by the Liaison Committee between the Central Norway Regional Health Authority and the Norwegian University of Science and Technology, and the Norwegian National Advisory Unit for functional MRI. The authors would like to thank senior neuroradiologist Kjell Arne Kvistad and Jana Rydland for performing the WMH scoring.

*LIFE Adult*

LIFE Adult is funded by the Leipzig Research Center for Civilization Diseases (LIFE). LIFE is an organizational unit affiliated to the Medical Faculty of the University of Leipzig. LIFE is funded by means of the European Union, by the European Regional Development Fund (ERDF) and by funds of the Free State of Saxony within the framework of the excellence initiative. We thank the participants of LIFE-Adult very much for their time and blood samples. We thank Kerstin Wirkner very much for running the LIFE study center.

*Lothian Birth Cohort 1936 (LBC1936), Scotland*

We thank the LBC1936 cohort members and research team staff who were involved in phenotypic and genotypic data collection, processing and preparation. The LBC1936 is supported by Age UK (Disconnected Mind programme grant). The work was undertaken by The University of Edinburgh Centre for Cognitive Ageing and Cognitive Epidemiology, part of the cross council Lifelong Health and Wellbeing Initiative (MR/K026992/1). The brain imaging was performed in the Brain Research Imaging Centre (www.bric.ed.ac.uk) a centre in the SINAPSE Collaboration (www.sinapse.ac.uk) supported by the Scottish Funding Council and Chief Scientist Office. Funding from the UK Biotechnology and Biological Sciences Research Council (BBSRC) and the UK Medical Research Council (MRC) is gratefully acknowledged. Genotyping was supported by a grant from the BBSRC (Ref. BB/F019394/1)

*Northern Manhattan Study (NOMAS)*

We gratefully acknowledge and thank the NOMAS participants. Funding was awarded through grants from the National Institute of Neurological Disorders and Stroke (R01 NS 29993) and the Evelyn F. McKnight Brain Institute.

*Older Australian Twins Study (OATS)*

We gratefully acknowledge and thank the OATS participants, their supporters and the OATS Research Team (current and former staff and students). This research was facilitated through Twins Research Australia, a national resource in part supported by a Centre for Research Excellence from the NHMRC. Funding for this study was awarded by the Australian National Health and Medical Research Council (NHMRC)/Australian Research Council Strategic Award (Grant 401162) and the NHMRC Project grant 1405325.

*PROspective Study of Pravastatin in the Elderly at Risk (PROSPER)*

The PROSPER study was supported by an investigator initiated grant obtained from Bristol-Myers Squibb. Prof. Dr. J. W. Jukema is an Established Clinical Investigator of the Netherlands Heart Foundation (grant 2001 D 032). Support for genotyping was provided by the seventh framework program of the European commission (grant 223004) and by the Netherlands Genomics Initiative (Netherlands Consortium for Healthy Aging grant 050-060-810).

*Rotterdam Study (RS)*

The generation and management of GWAS genotype data for the Rotterdam Study (RS I, RS II, RS III) was executed by the Human Genotyping Facility of the Genetic Laboratory of the Department of Internal Medicine, Erasmus MC, Rotterdam, The Netherlands. The GWAS datasets are supported by the Netherlands Organisation of Scientific Research NWO Investments (nr. 175.010.2005.011, 911-03-012), the Genetic Laboratory of the Department of Internal Medicine, Erasmus MC, the Research Institute for Diseases in the Elderly (014-93-015; RIDE2), the Netherlands Genomics Initiative (NGI)/Netherlands Organisation for Scientific Research (NWO) Netherlands Consortium for Healthy Aging (NCHA), project nr. 050-060-810. We thank Pascal Arp, Mila Jhamai, Marijn Verkerk, Lizbeth Herrera and Marjolein Peters, MSc, and Carolina Medina-Gomez, MSc, for their help in creating the GWAS database, and Karol Estrada, PhD, Yurii Aulchenko, PhD, and Carolina Medina-Gomez, MSc, for the creation and analysis of imputed data.

The Rotterdam Study is funded by Erasmus Medical Center and Erasmus University, Rotterdam, Netherlands Organization for the Health Research and Development (ZonMw), the Research Institute for Diseases in the Elderly (RIDE), the Ministry of Education, Culture and Science, the Ministry for Health, Welfare and Sports, the European Commission (DG XII), and the Municipality of Rotterdam. HHHA was supported by ZonMW grant number 916.19.151. The authors are grateful to the study participants, the staff from the Rotterdam Study and the participating general practitioners and pharmacists.

*Sydney Memory and Ageing Study (Sydney MAS*

We gratefully acknowledge and thank the Sydney MAS participants, their supporters and the Sydney MAS Research Team (current and former staff and students). Funding was awarded from the Australian National Health and Medical Research Council (NHMRC) Program Grants (350833, 568969, 109308).

*Three-City Dijon (3C-Dijon)*

The Three City (3C) Study is conducted under a partnership agreement among the Institut National de la Santé et de la Recherche Médicale (INSERM), the University of Bordeaux, and Sanofi-Aventis. The Fondation pour la Recherche Médicale funded the preparation and initiation of the study. The 3C Study is also supported by the Caisse Nationale Maladie des Travailleurs Salariés, Direction Générale de la Santé, Mutuelle Générale de l’Education Nationale (MGEN), Institut de la Longévité, Conseils Régionaux of Aquitaine and Bourgogne, Fondation de France, and Ministry of Research–INSERM Programme “Cohortes et collections de données biologiques. Christophe Tzourio and Stéphanie Debette have received investigator-initiated research funding from the French National Research Agency (ANR) and from the Fondation Leducq. Stéphanie Debette is also supported by a starting grant from the European Research Council, a grant from the Joint Programme of Neurodegenerative Disease research, and the Initiative of Excellence of Bordeaux University. We thank Dr. Anne Boland (CNG) for her technical help in preparing the DNA samples for analyses. This work was supported by the National Foundation for Alzheimer’s disease and related disorders, the Institut Pasteur de Lille, the labex DISTALZ and the Centre National de Génotypage.

*United Kingdom Biobank (UKB)*

We gratefully acknowledge the UK Biobank for making the data available, and to all UKB participants who generously participated in this study to make this resource possible. F.A.-A. acknowledges funding from the UK Medical Research Council and the Wellcome Trust via UK Biobank. S.M.S. receives further support from the Wellcome Trust. J.M. acknowledges funding for this work from the European Research Council (ERC; grant 617306) and the Leverhulme Trust. M.J. is supported by the National Institute for Health Research (NIHR) Oxford Biomedical Research Centre (BRC). L.G. is supported by the MRC Dementias Platform UK and the Monument Trust Discovery Award from Parkinson’s UK (Oxford Parkinson’s Disease Centre).

*Vietnam Era Twin Study of Aging (VETSA)*

This work was supported by grants from the U.S. National Institute on Aging (R01 AG050595, R01 AG022381, R03 AG046413, K08 AG047903, P01 AG055367, R01 AG059329) and the VA San Diego Center of Excellence for Stress and Mental Health. The content is the responsibility of the authors and does not necessarily represent official views of the NIA, NIH, or VA. The Cooperative Studies Program of the U.S. Department of Veterans Affairs provided financial support for development and maintenance of the Vietnam Era Twin Registry. We would also like to acknowledge the continued cooperation and participation of the members of the VET Registry and their families.

3. **Supplementary Figures**

*Supplementary Figure 1*. Regional Plots of genome-wide significant SNPs for periventricular WMHs

| 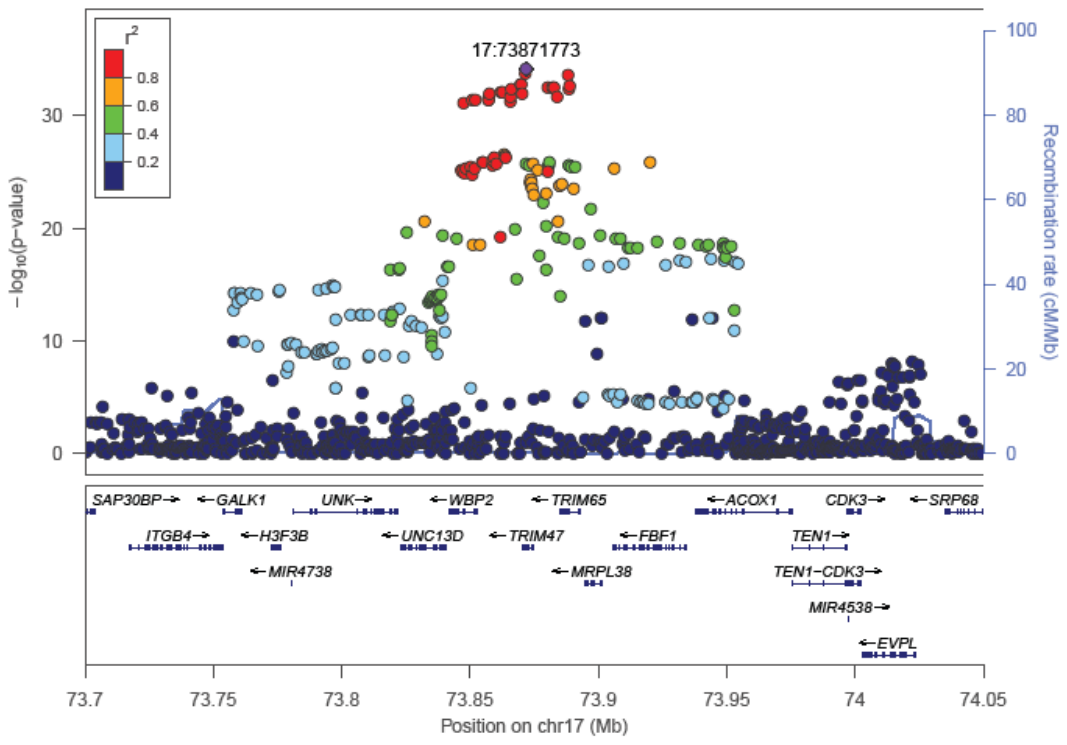 | 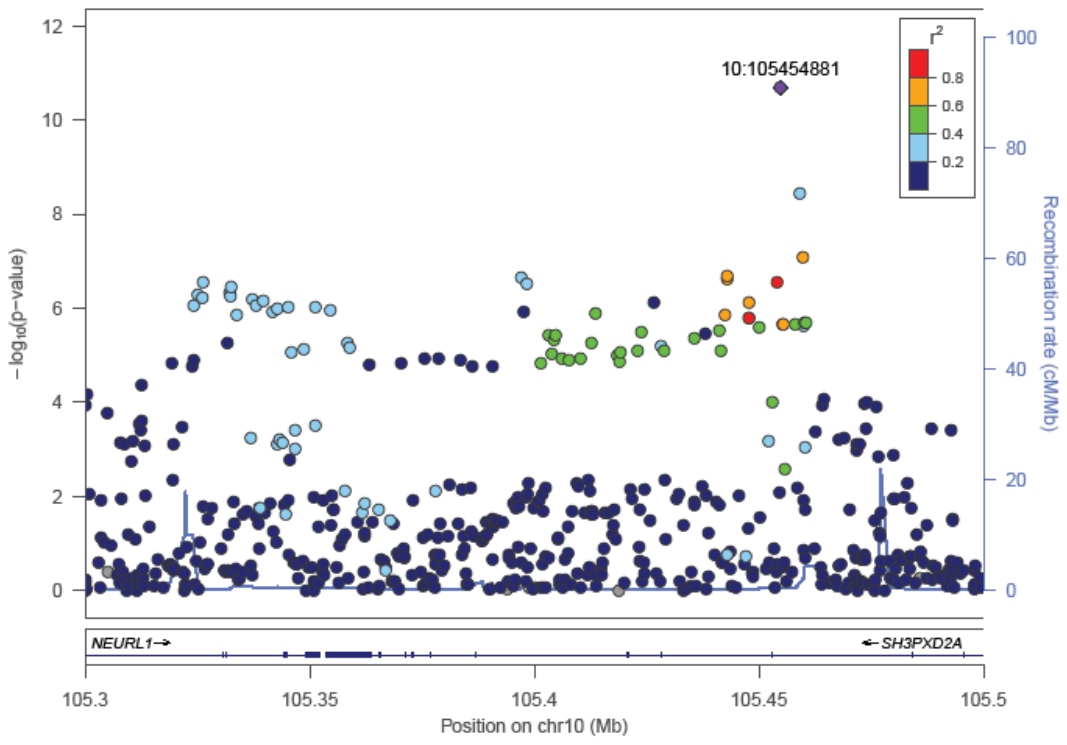 | 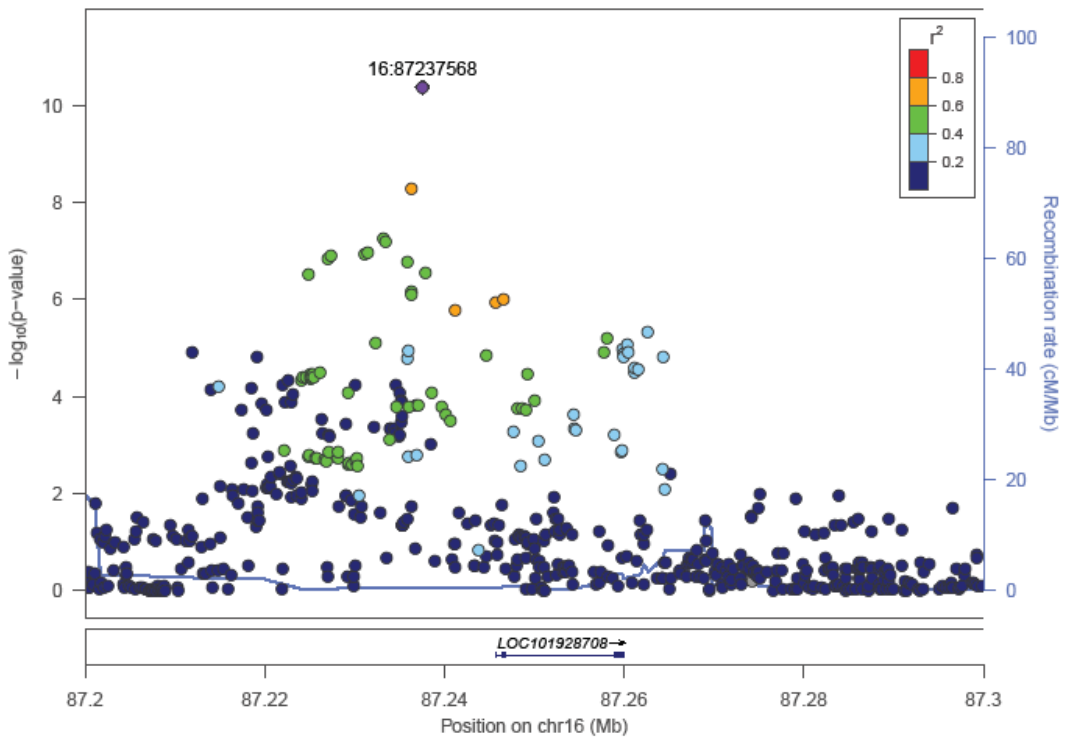 |
| --- | --- | --- |
| (a) Chr 17q25.1 | (b) Chr10q24.33 | (c) Chr16q24.2 |
| 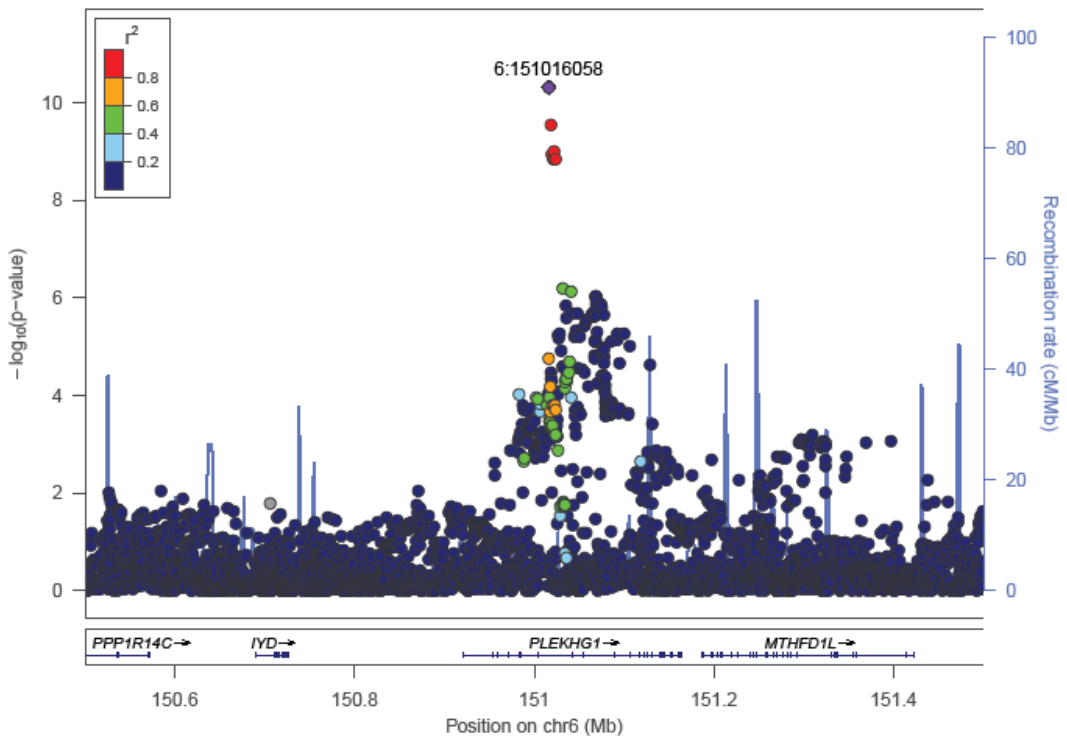 | 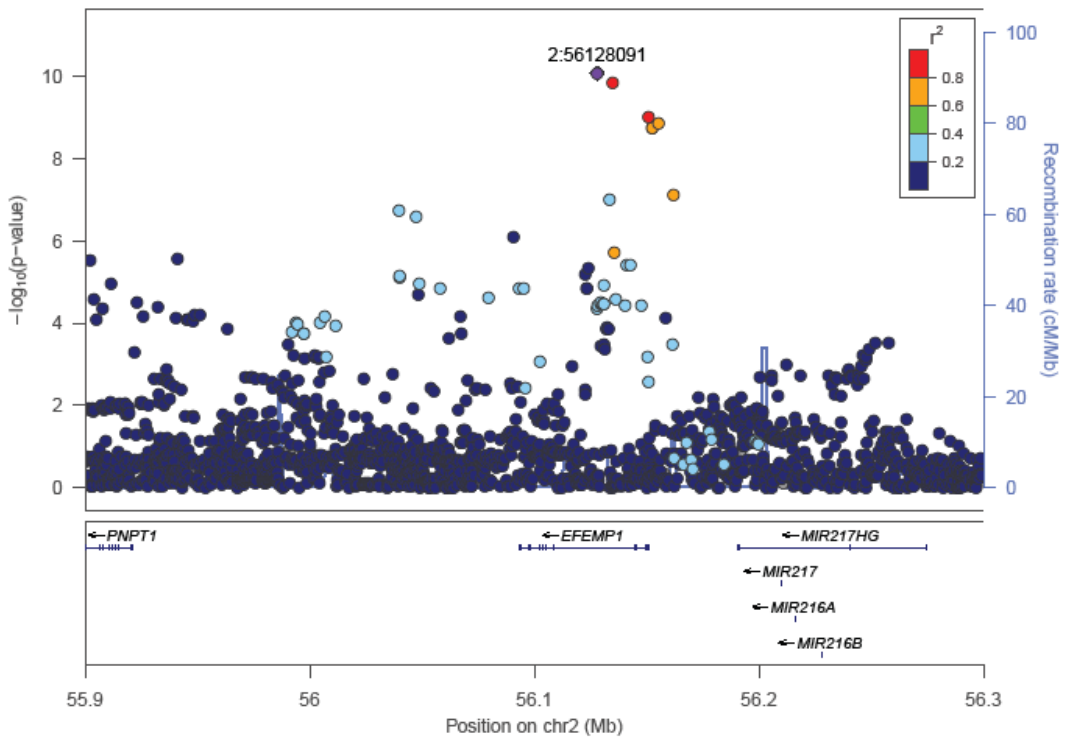 | 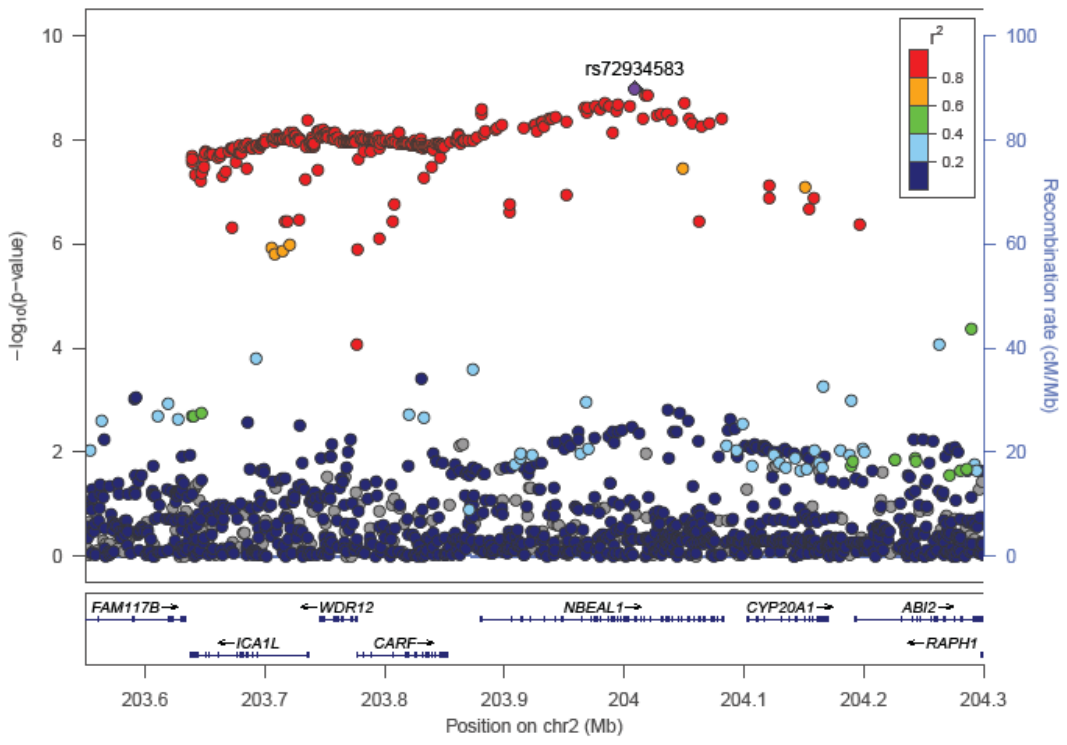 |
| (d) Chr6q25.1 | (e) Chr2p16.1 | (f) Chr2q33.2 |
| 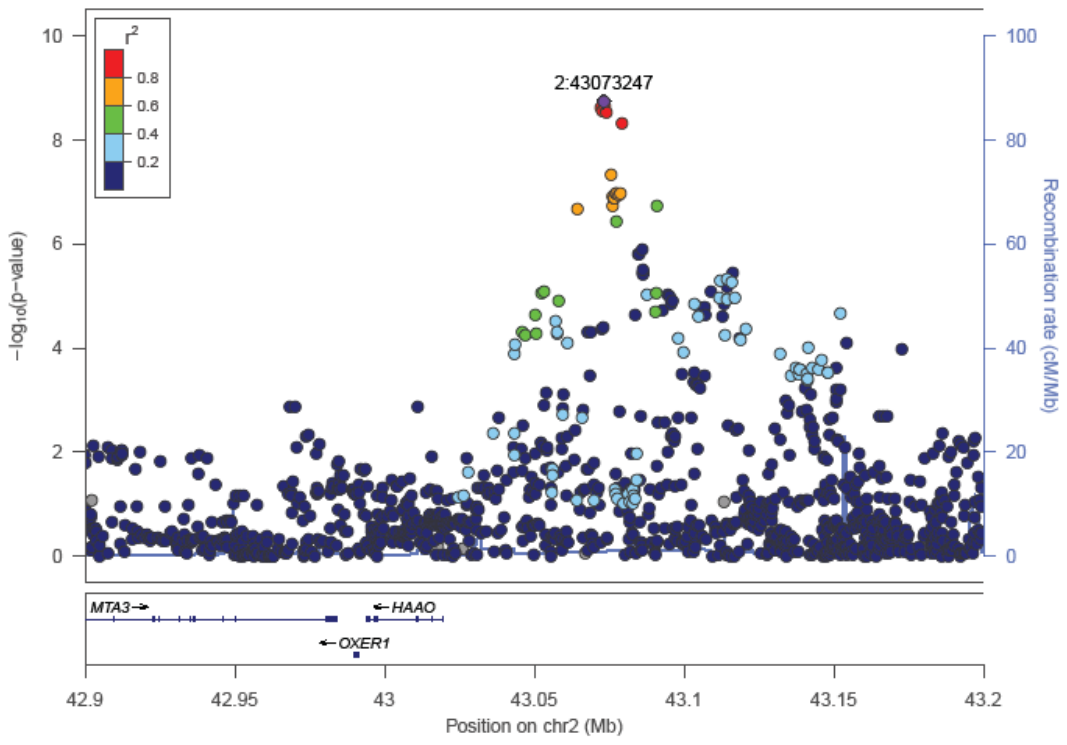 | 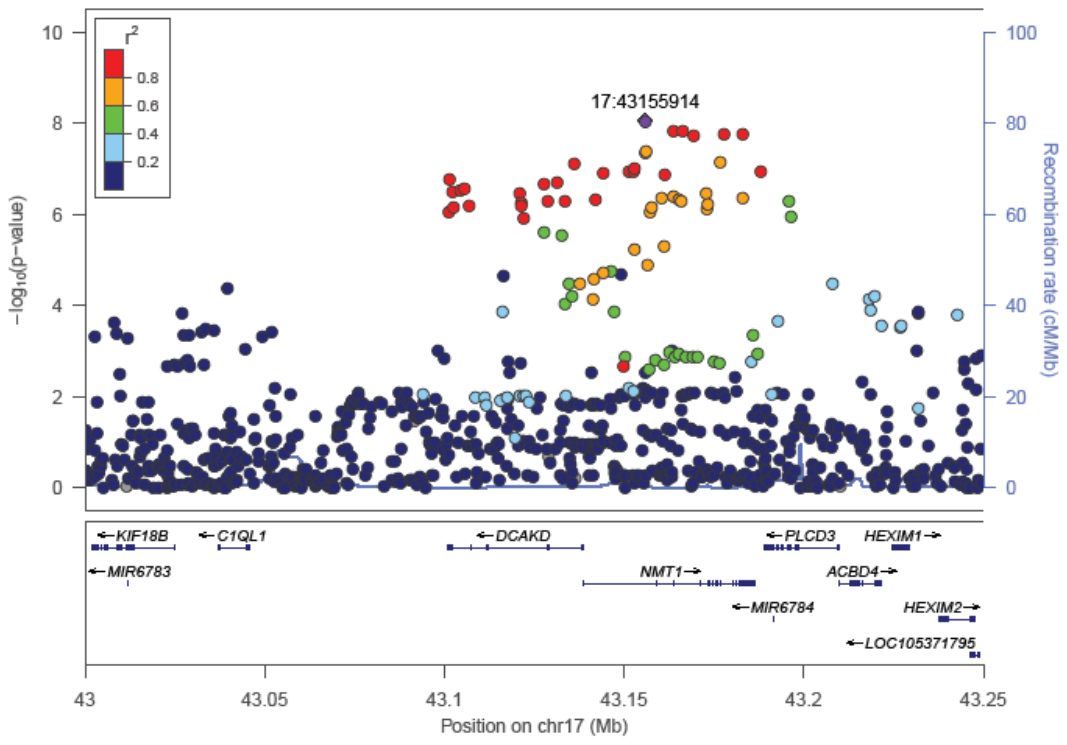 | 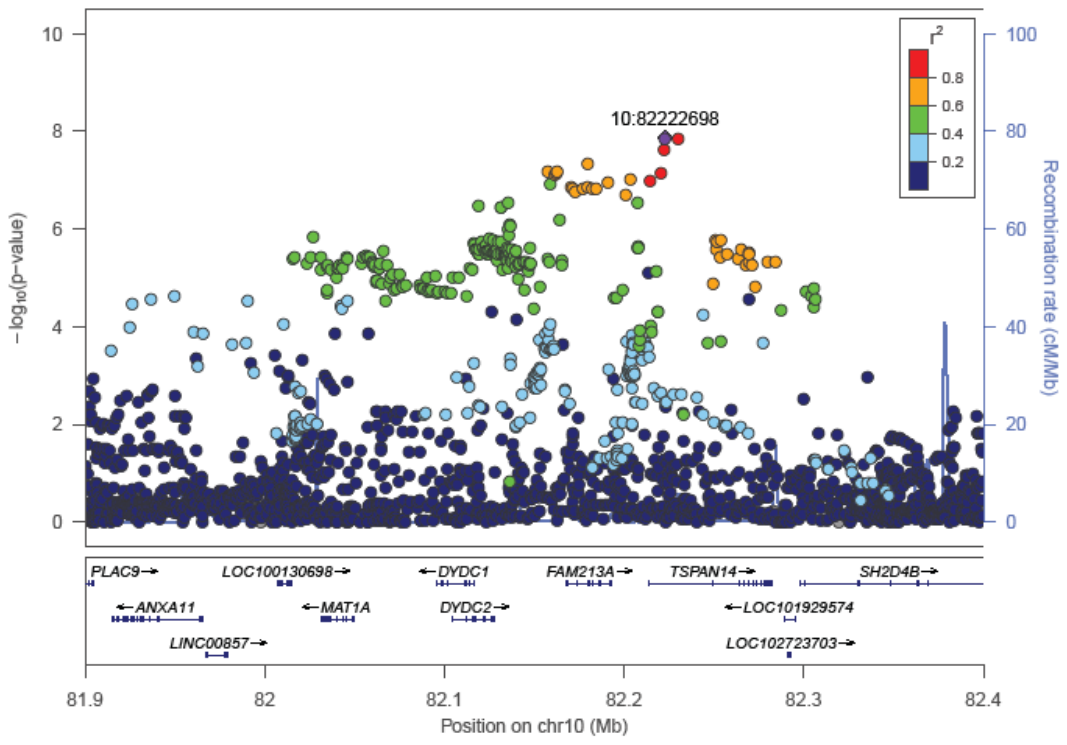 |
| (g) Chr2p21 | (h) Chr17q23.1 | (i) Chr10q23.1 |
| 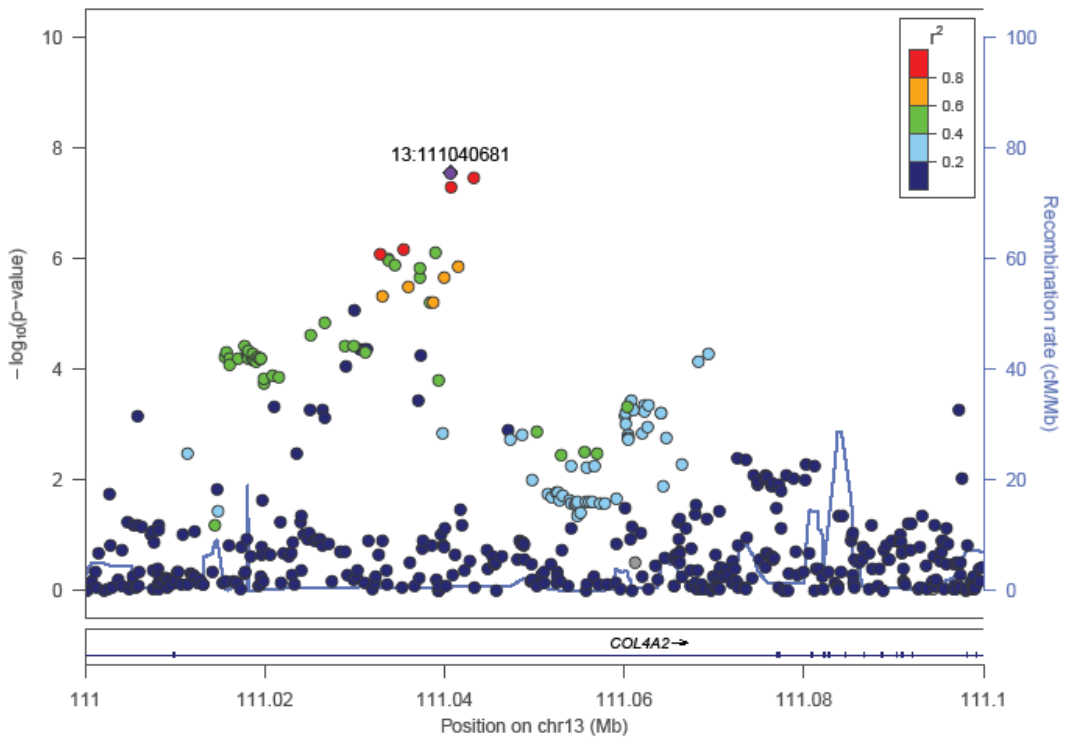 | 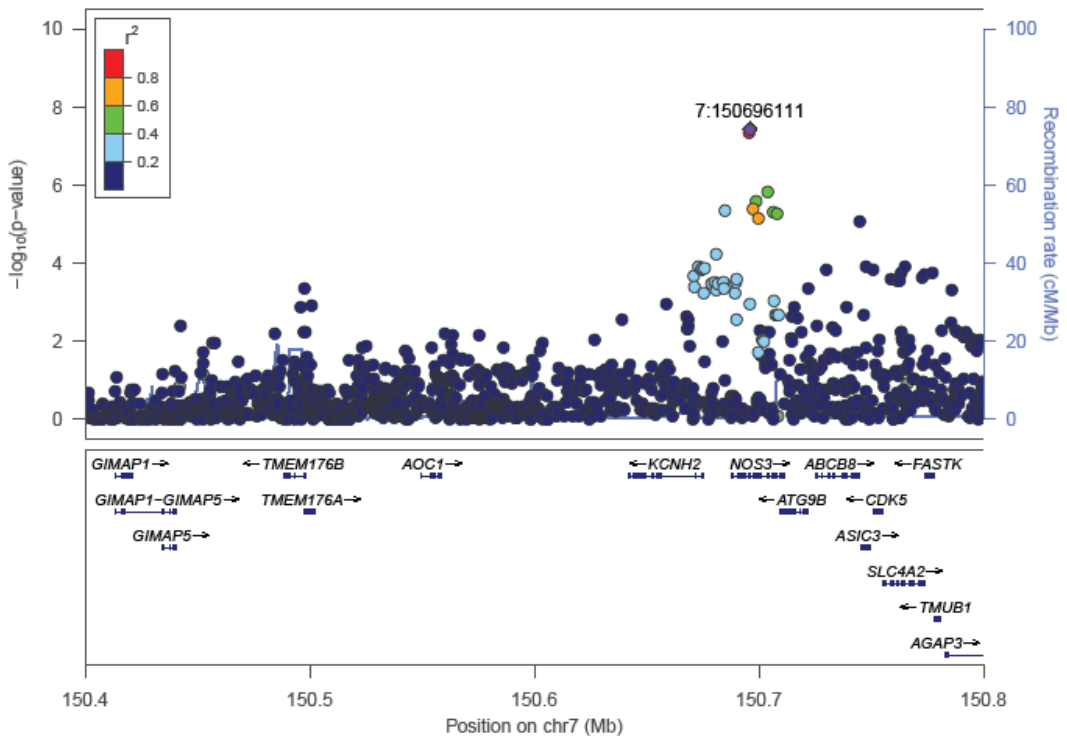 |  |
| (j) Chr13q34 | (k) Chr7q36.1 |  |

***Supplementary Figure 2.*** Circos plots for each chromosome containing significant loci. Outer ring shows SNPs <.05 with the most significant SNPs located towards the outermost ring. SNPs in high LD with the independent significant SNPs in each locus are colored in red (r^2^>.8)-blue (r^2^>.2); no LD (grey). Genomic risk loci are colored in dark blue (2^nd^ layer). Genes are mapped by chromatin interaction (orange), eQTL (green) or both (red).

| 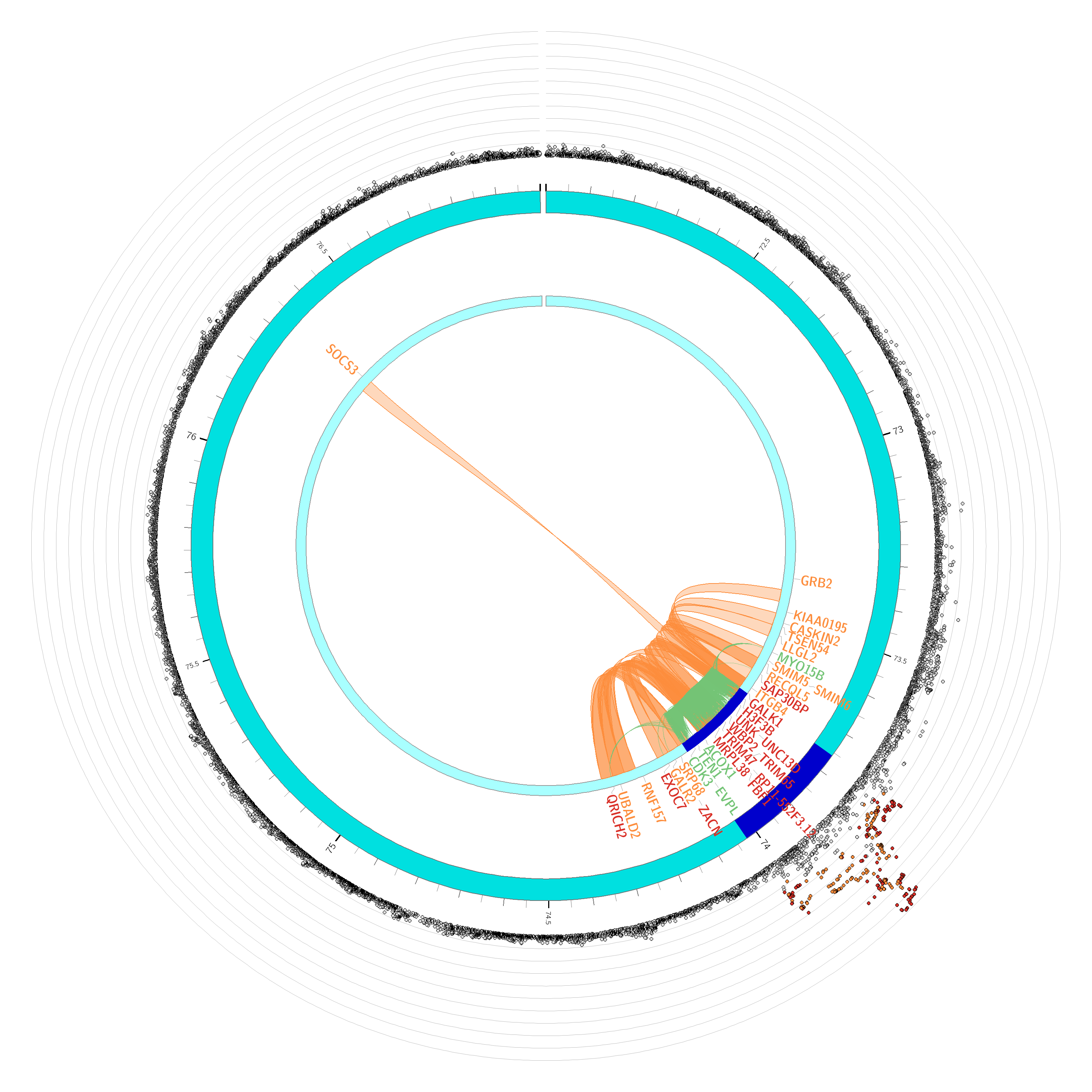 | 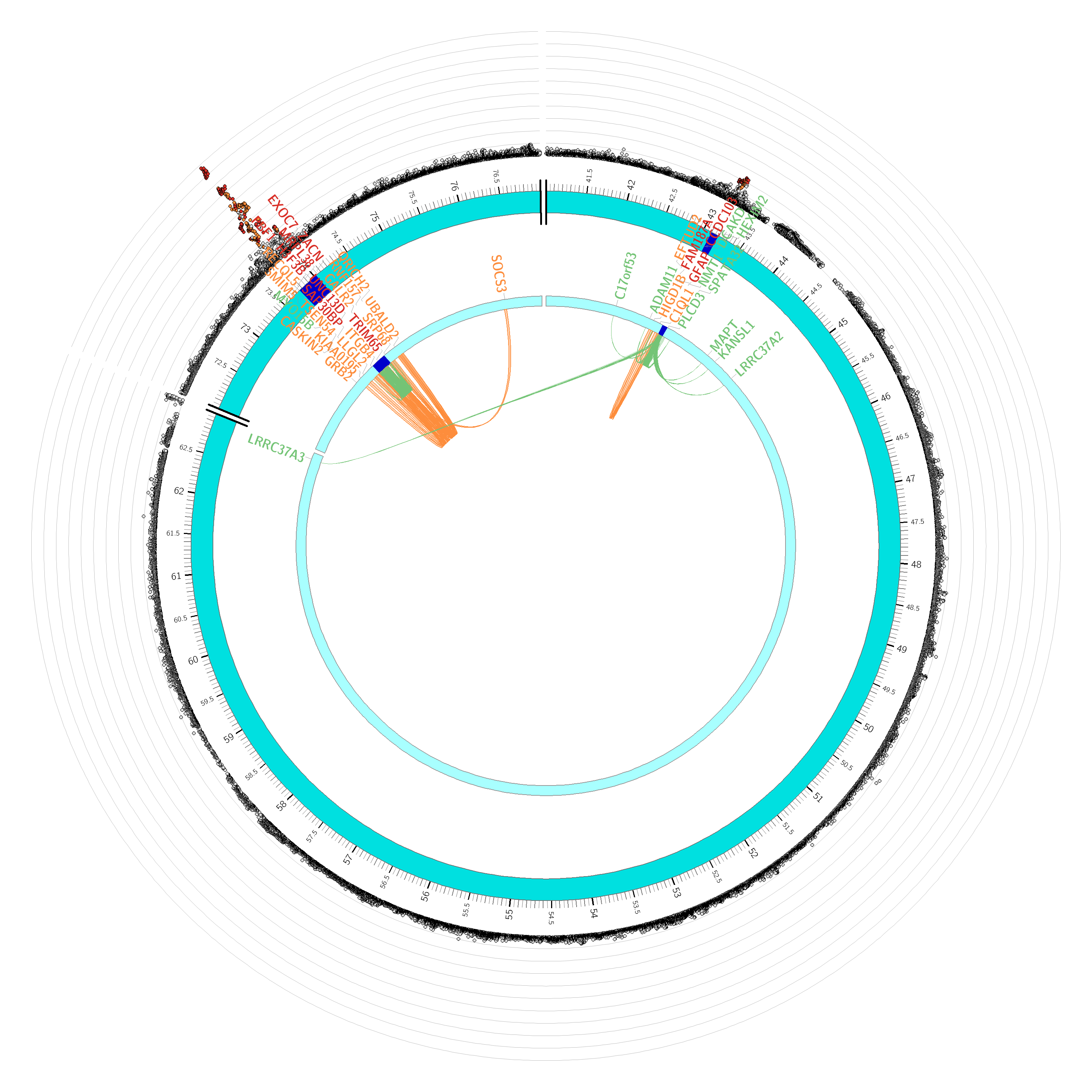 |
| --- | --- |
| Deep Chr 17 | Periventricular Chr 17 |
| 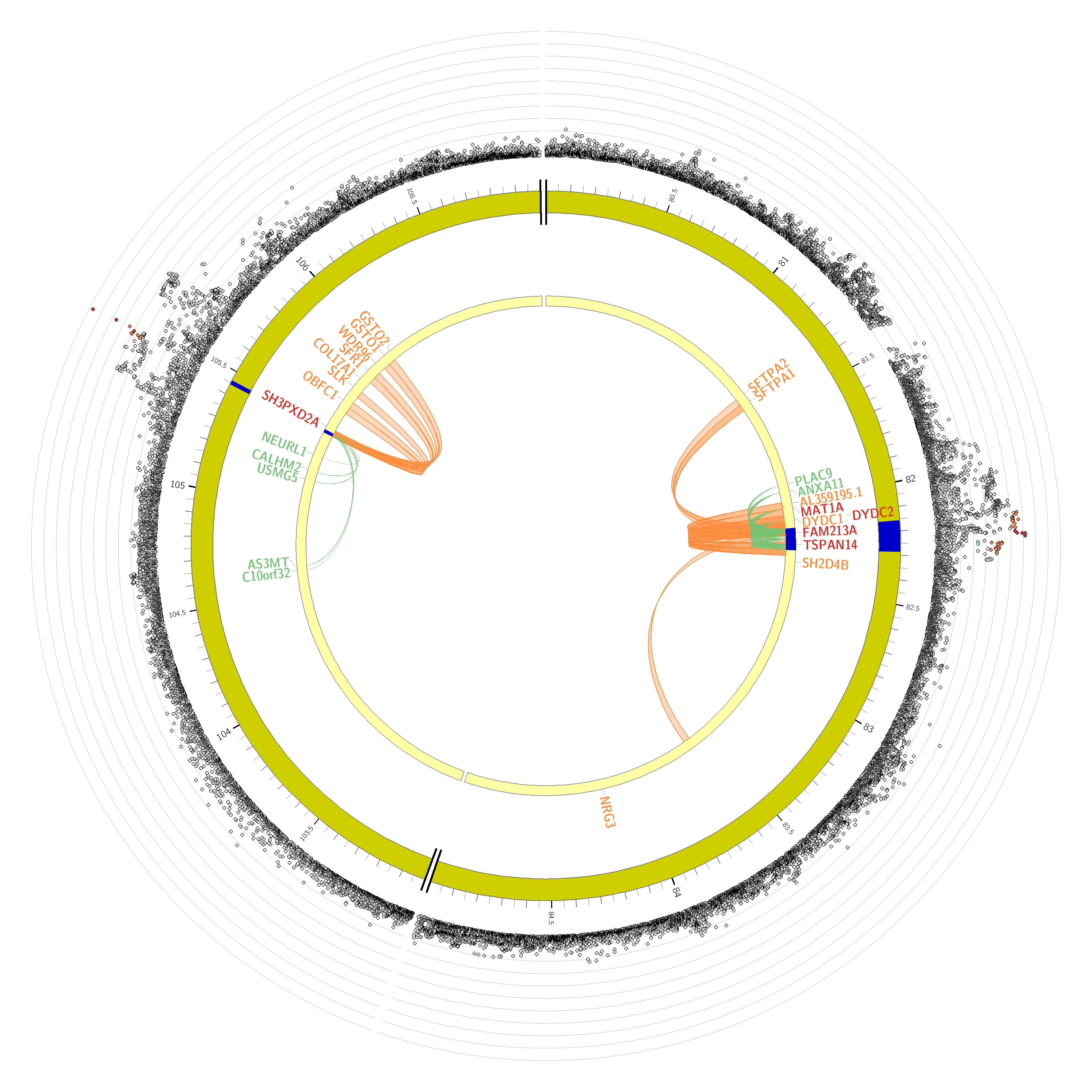 | 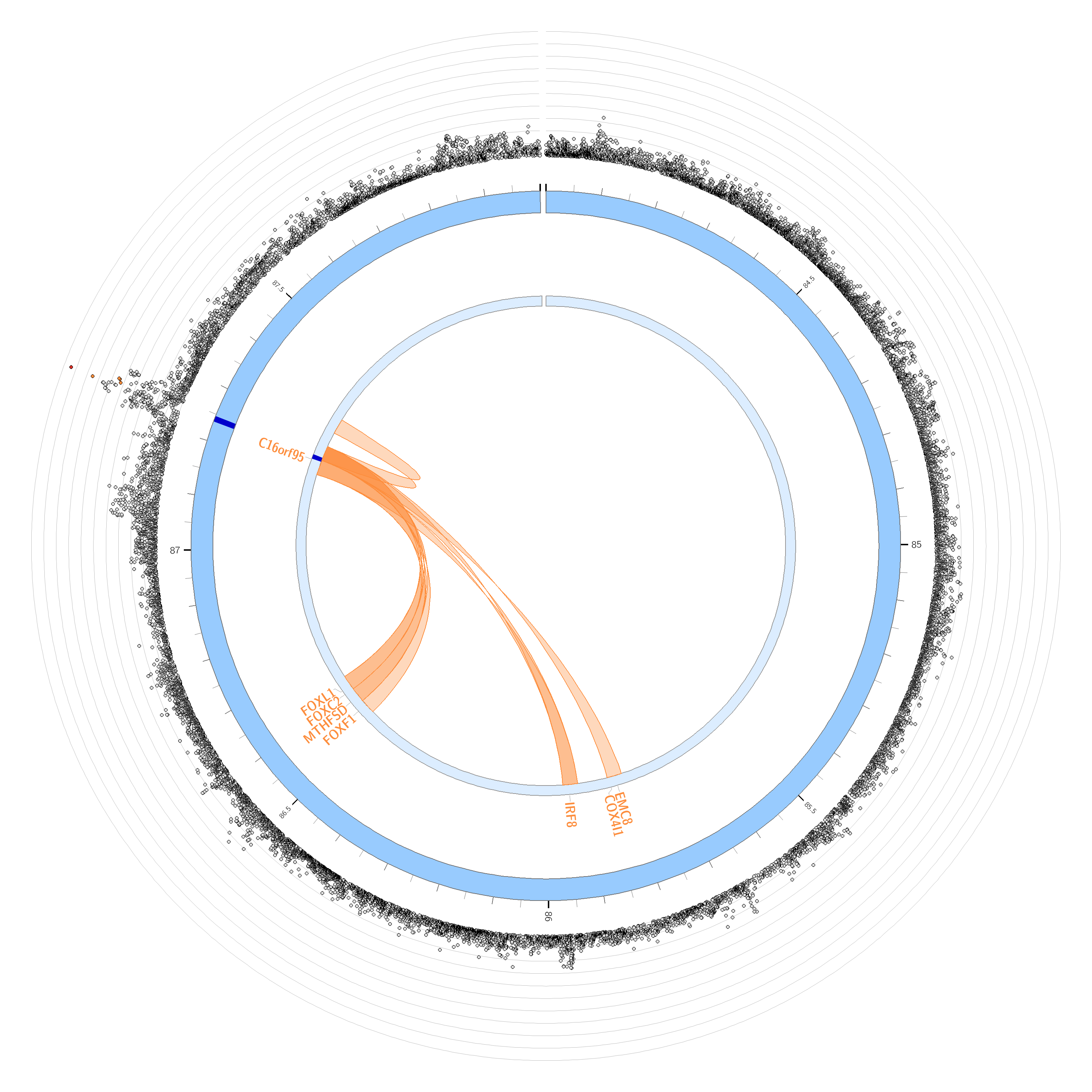 |
| Periventricular Chr10 | Periventricular Chr16 |
| 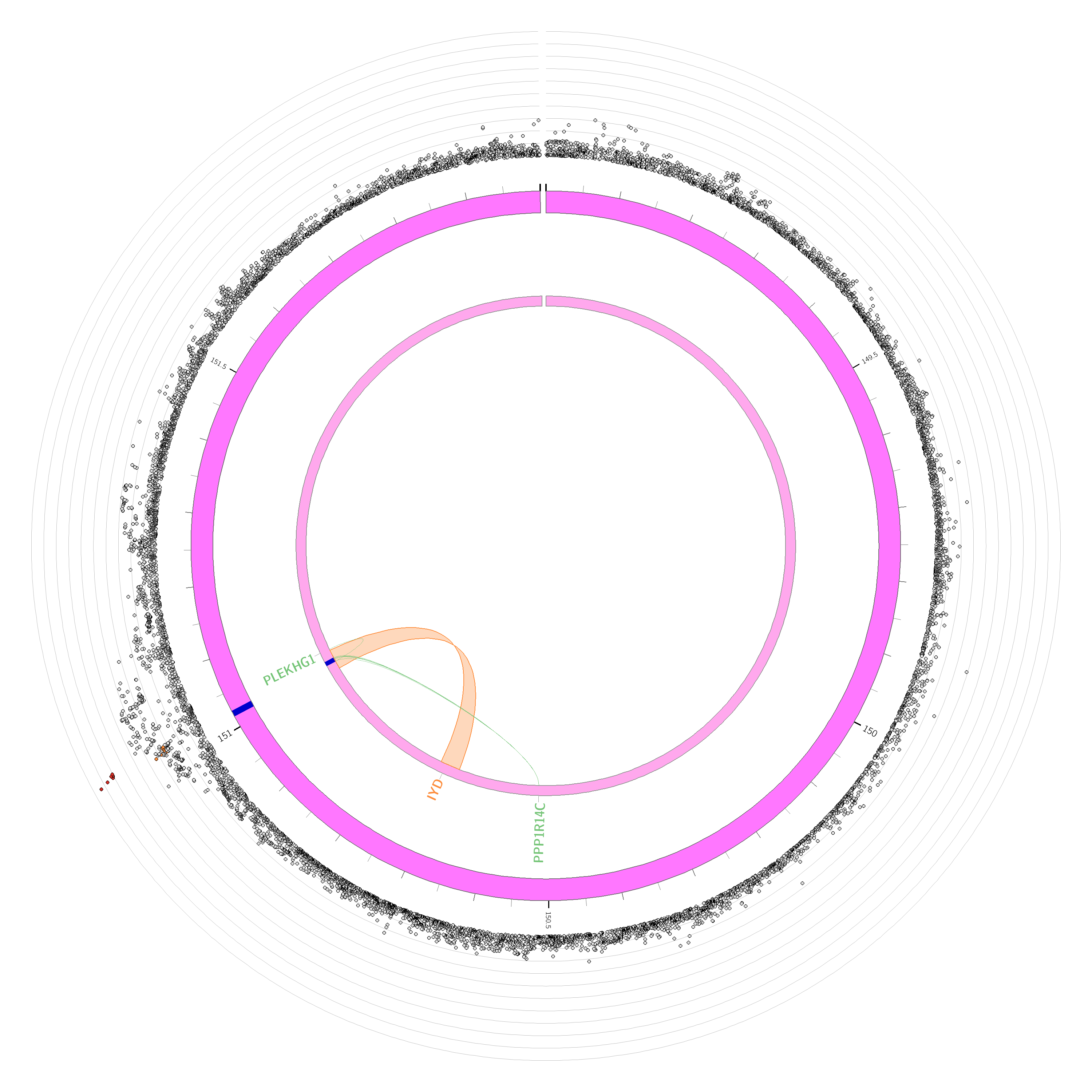 | 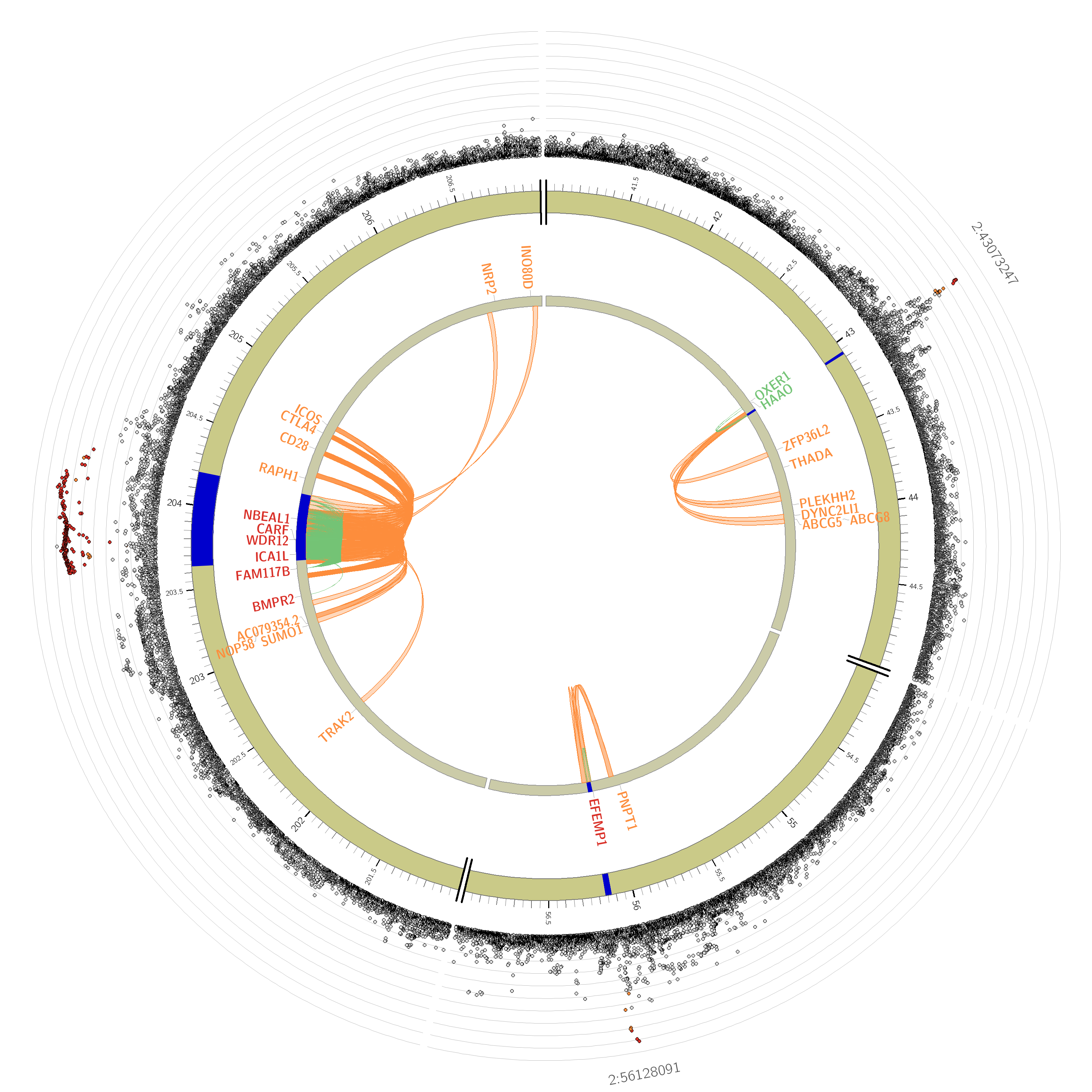 |
| Periventricular Chr6 | Periventricular Chr2 |
| 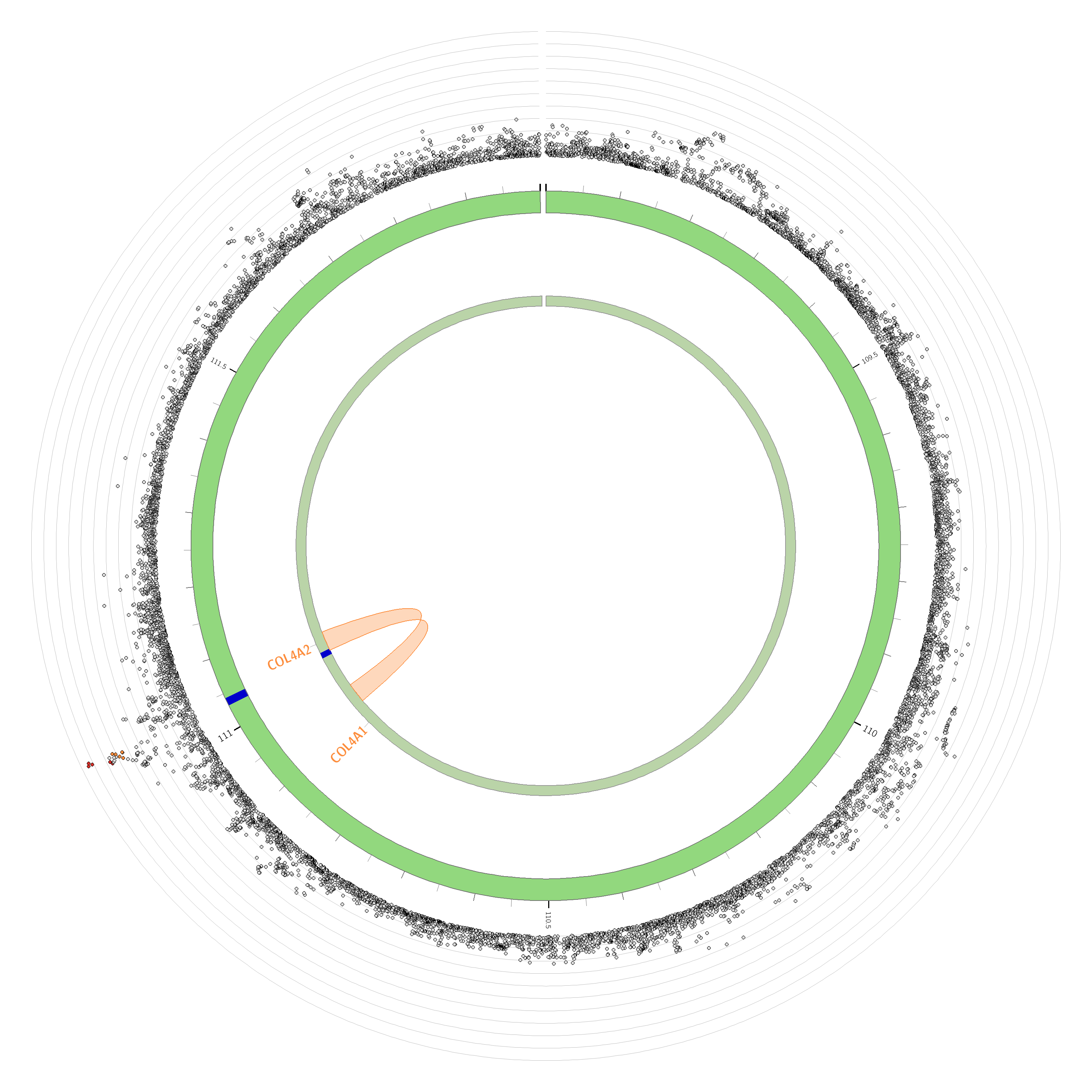 | 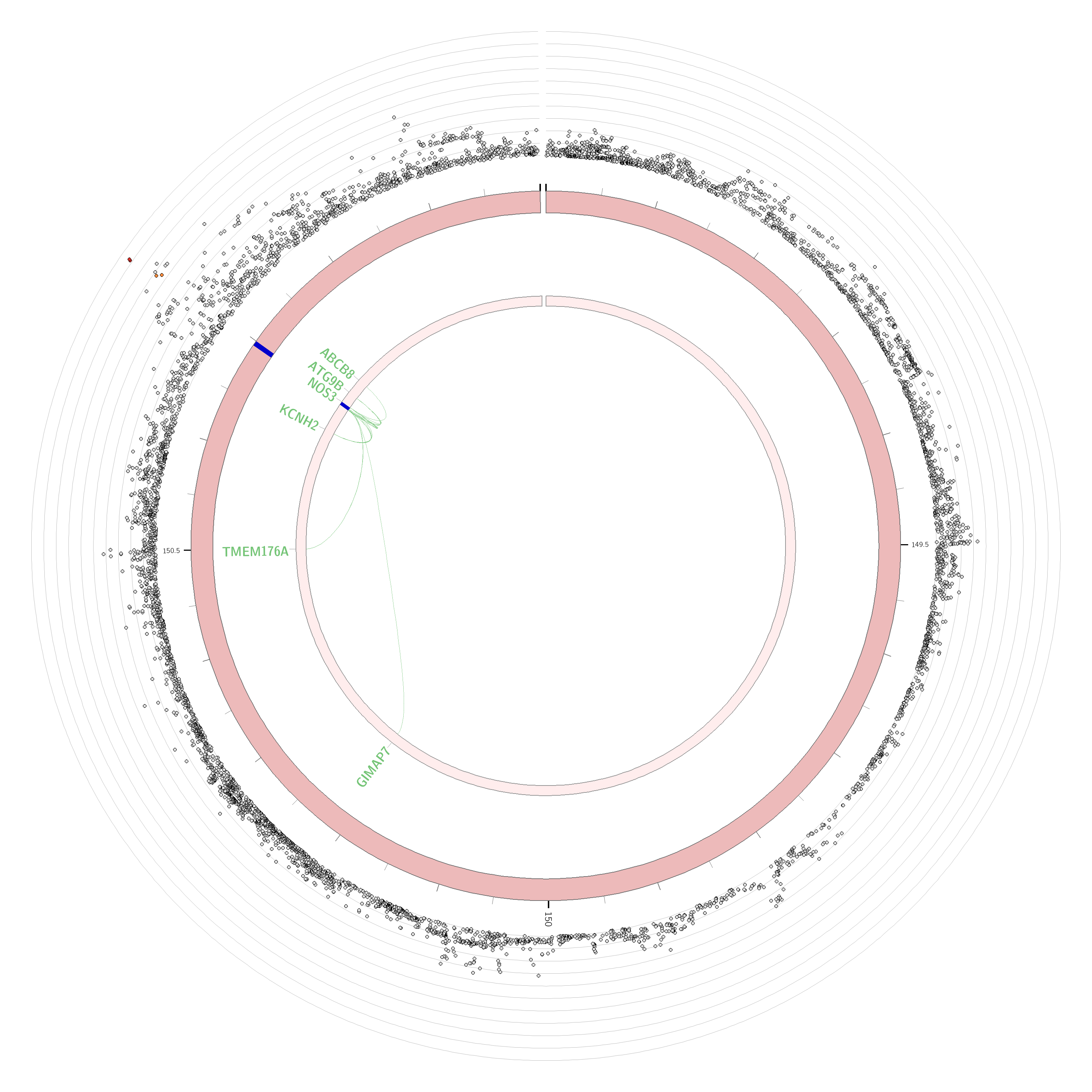 |
| Periventricular Chr13 | Periventricular Chr7 |

4. **Supplementary Table Legends**

| **Supplementary Table** | **Legend** |
| --- | --- |
| **1** | Individual study demographics |
| **2** | Study MRI and WMH extraction details |
| **3** | Study genotyping, imputation and association analysis details |
| **4** | Discovery *deep* white matter hyperintensity results for Model 1, genome-wide significant (p<5e-8, n=167) and suggestive results (P<1e-5, n=173). n= 340 SNPs. |
| **5** | Discovery *periventricular* white matter hyperintensity results for Model 1, genome-wide significant (p<5e-8, n=217) and suggestive results (p<1e-5, n=882), n= 1099 SNPs. |
| **6** | Replication of Discovery Model 1 for DWMH by the UKB. 139 out of 167 genome-wide significant SNPs were available for replication in the UKB. 127/139 were significant (Bonferroni adjusted p-value of .05/139 = 3.6e-4) and in the correct direction. Twelve were not significant, 11/12 were however, in the correct direction |
| **7** | Replication of genome-wide significant results for PVWMH discovery Model 1 by the UKB. 181/217 of Model 1 genome-wide significant SNPs were available in the UKB. Using a Bonferroni adjusted p-value for replication of 2.76e-4 (.05/181), 162/181 were significant and were all in the correct direction. Of the 19 that were not significant, only 2 were not in the correct direction. |
| **8** | Discovery DWMH results for Model 2, genome-wide significant (p<5e-8, n=166) and suggestive results (p<1e-5, n=175). Total n=341 SNPs. |
| **9** | Discovery PVWMH results for Model 2, genome-wide significant (p<5e-8, n=200) and suggestive results (p<1e-5, n=825. Total n = 1025 SNPs |
| **10** | DWMH Phase II Analysis results (discovery & replication UKB cohorts). 381 genome-wide (p<=5e-8, n=236) and suggestive (p>=5e-8 & <1e-5, n=145) SNPs. Notes. The allele frequency, the effect size (beta) and standard error are given with reference to allele 1. Effect sizes are additive effects for each copy of allele 1. |
| **11** | PVWMH Phase II Analysis results (discovery & replication UKB cohorts). 1632 genome-wide (p<=5e-8, n=513) and suggestive (p>=5e-8 & <1e-5, n=1119) SNPs. Notes. The allele frequency, the effect size (beta) and standard error are given with reference to allele 1. Effect sizes are additive effects for each copy of allele 1 |
| **12** | GWAS catalogue results for DWMH SNPs using the Phase II meta-analysis results |
| **13** | GWAS catalogue results for PVWMH SNPs using the Phase II meta-analysis results |
| **14** | DWMH gene-based analysis (MAGMA) identified 13 significant genes (Bonferroni p-value =.05/18745=2.66e-6). Highlighted genes are significant after Bonferroni correction |
| **15** | PVWMH gene-based analysis (MAGMA) identified 32 significant genes (Bonferroni p-value =.05/18745=2.66e-6). Highlighted genes are significant after Bonferroni correction |
| **16** | Genetic correlations with DWMH (Phase II), PVWMH (Phase II), Alzheimer's disease (Lambert et al., 2013) and stroke phenotypes (Malik et al., 2018, Woo et al., 2014) |
| **17** | Genetic overlap between PVWMH, DWMH and stroke types |
| **18** | Replication of previously identified total WMH SNPs from GWAS in DWMH and PVWMH. Highlighted p-values are significant after Bonferroni correction (.05/48= 0.00104) |
